# Piezo1 ion channels are capable of conformational signaling

**DOI:** 10.1101/2024.05.28.596257

**Authors:** Amanda H. Lewis, Marie E. Cronin, Jörg Grandl

## Abstract

Piezo1 is a mechanically activated ion channel that senses forces with short latency and high sensitivity. Piezos undergo large conformational changes, induce far-reaching deformation onto the membrane, and modulate the function of two-pore potassium (K2P) channels. Taken together, this led us to hypothesize that Piezos may be able to signal their conformational state to other nearby proteins. Here, we use chemical control to acutely restrict Piezo1 conformational flexibility and show that Piezo1 conformational changes, but not ion permeation through it, are required for modulating the K2P channel TREK1. Super-resolution imaging and stochastic simulations further reveal that both channels do not co-localize, which implies that modulation is not mediated through direct binding interactions; however, at high Piezo1 densities, most TREK1 channels are within the predicted Piezo1 membrane footprint, suggesting the footprint may underlie conformational signaling. We speculate that physiological roles originally attributed to Piezo1 ionotropic function could, alternatively, involve conformational signaling.

## Introduction

Force-gated ion channels (FGICs) have evolved to transduce mechanical force into ion flux with high speed and sensitivity. For example, Piezo1 is activated rapidly (within ∼1 ms) and directly by membrane tension with a tension of half-maximal activation (T_50_) of only 1.4 mN/m, making it the most sensitive of all currently known FGICs^1,2^. This exquisite speed and sensitivity underlie its key role in many physiological processes involving the transduction of both external forces, such as mechanical itch, and internal forces, such as cellular stiffness^3–5^.

In the canonical mechanism of Piezo1 function, membrane tension induces conformational changes that include the opening of its central pore, resulting in a flux of cations that relay a mechanical force into a cellular signal. For example, Piezo1-mediated calcium flux activates the Gardos channel in red blood cells and modulates transcription pathways in many cell types^6–9^. Likewise, excitatory currents through the closely related isoform Piezo2 are required for action potential firing in mechanosensory afferents that mediate the sense of light touch^10,11^. The existence of Piezo1 and Piezo2 splice variants with altered unitary conductance and calcium permeability further hints at the physiological importance of their ionotropic activity^12,13^.

Some ion channels, however, can also directly modulate other proteins through flux-independent *‘conformational signaling’*. For example, the voltage-gated calcium channel Ca_v_1.2 requires both voltage-dependent conformational changes and calcium influx to drive neuronal gene expression^14,15^. Similarly, NMDA receptors undergo ligand-induced conformational changes that are sufficient to mediate structural plasticity of dendritic spines, even in the complete absence of ion permeation^16–18^.

The ability of Piezo1 ion channels to influence other proteins via conformational signaling has not been directly tested, but is plausible for several reasons: First, Piezo ion channels are exceptionally large proteins, with a diameter of ∼28 nm, extreme intrinsic curvature, and substantial conformational flexibility^19,20^. Upon mechanical force, the bowl-like channel flattens, resulting in a displacement of its propeller-like blades by ∼10 nm, a large movement compared to conformational changes in most other ion channels^21–23^. Second, in its curved conformation, Piezo1 deforms its surrounding membrane bilayer, giving rise to a large, curved membrane footprint that extends far beyond the protein itself^19,24–26^. Flattening of the Piezo protein necessarily reduces the curvature of its membrane footprint, potentially eliminating it entirely. Together, these membrane-associated changes and the large diameter of Piezo1 mean that its conformation may manifest itself 10s of nm away from its central pore. Third, both Piezo1 and Piezo2 channels have been shown to substantially potentiate the activity of force-gated two-pore potassium (K2P) channels, including TREK1^27^. The mechanism of this modulation is unknown, but conformational signaling, potentially through Piezo’s large, flexible structure and/or its curved membrane footprint, is an enticing possibility, because this modulation persists under conditions of zero net ion flow through Piezo1. Finally, TREK1 modulation by Piezo1 occurs naturally in adipose stem cells and mouse gingival fibroblasts, where it also contributes to wound healing in a gum injury mouse model, indicating that conformational signaling may be physiologically relevant^27^. Given all this evidence, we recognized conformational signaling as an enticing mechanism and set out to test it systematically.

## Results

### Piezo ion channels potentiate and modulate TREK1 mechanosensitive currents

We hypothesized that Piezo1 ion channels are capable of conformational signaling. To test this, we turned to the intriguing finding by Glogowska and colleagues that Piezo1 potentiates TREK1, and first aimed to reproduce this finding and carefully validate our model system and protocols^27^. Specifically, we transfected Neuro2A cells lacking endogenous Piezo1 expression (Neuro2A-Piezo1ko) with mouse TREK1, mouse Piezo1, or both constructs combined, and recorded currents via cell-attached patch clamp electrophysiology. To isolate currents mediated by TREK1 and Piezo1, we recorded at holding potentials of -80 mV or 0 mV, which we validated are near their respective reversal potentials (**Figure S1**, see **Methods**). Mechanical sensitivity was probed by applying brief negative pressure steps through the patch pipette; for simplicity, in the main text we report current properties at -80 mmHg, where both TREK1 and Piezo1 currents are maximal. When expressing Piezo1 alone, outward currents at 0 mV were negligible (median current: 2.4 pA (1^st^-3^rd^ quartiles: 2.1 ー 4.1 pA, n=25) and when expressing TREK1 alone, inward currents at -80 mV were similarly small (-5.9 pA (-5.2 ー -7.1 pA, n=58); **Figure 1A-E**). Such minor current amplitudes likely result from seal noise as well as small changes in leak and capacitance during the pressure step^28,29^, and indirectly confirm that Neuro2A-Piezo1ko cells do not endogenously express Piezo or K2P channels and are a suitable background for our experiments.

**Figure 1.**
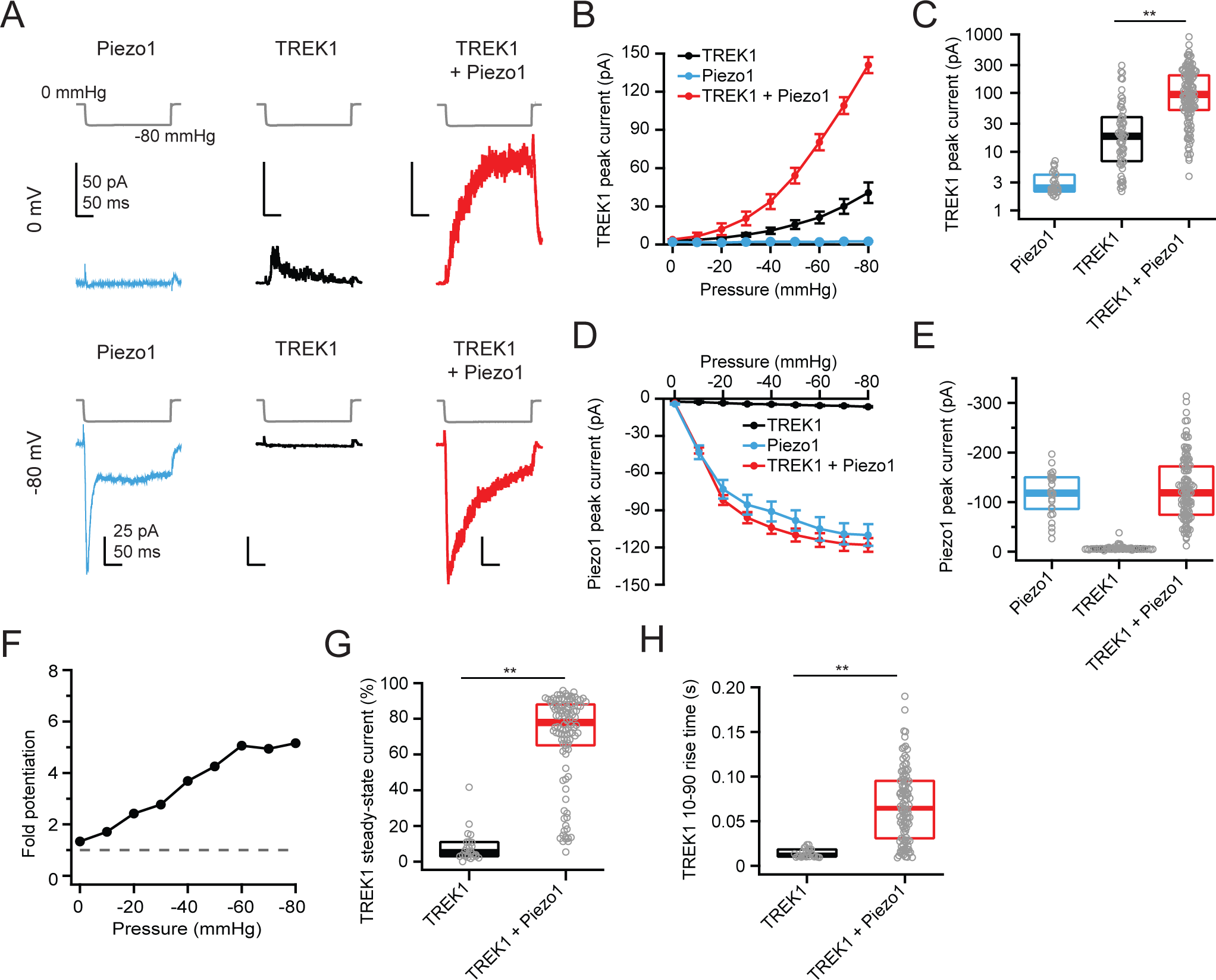
Piezo1 potentiates TREK1 peak and steady-state currents. **A**, Pressure protocol (gray) and currents from representative cell-attached patches from Neuro2A-Piezo1ko cells expressing Piezo1 (blue), TREK1 (black), or TREK1 + Piezo1 (red). Pressure step was to -80 mmHg and voltage was 0 mV (top) or -80 mV (bottom). **B**, Mean pressure-response curves at 0 mV, generated from full pressure step protocol. n=58 (TREK1), n=25 (Piezo1), n=121 (TREK1 + Piezo1). **C**, Box-and-whisker plot for peak outward current elicited by a pressure step to -80 mmHg at 0 mV. **D**, as in B, for inward currents at -80 mV. **E**, as in C, for currents at -80 mV **F**, Fold potentiation of TREK1 peak currents as a function of pressure. **G**, TREK1 steady-state current at -80 mmHg for cells expressing TREK1 (black, n=24) or TREK1 + Piezo1 (red, n=106). **H**, TREK1 10-90% rise time at 0 mV for cells expressing TREK1 (black, n=24) or TREK1 + Piezo1 (red, n=106). Significance assessed with a Mann-Whitney U test; ∗∗p < 0.0001. Error bars are ±SEM.

Co-expression of Piezo1 strongly modulates TREK1 currents in several ways^27^. First, TREK1 peak currents increased from 18.3 pA (7.0 ー 38.9 pA, n=58) in the absence of Piezo1 to 94.3 pA (51.4 ー 200.0 pA, n=121) in the presence of Piezo1, a more than 5-fold increase (**Figure 1B-C,F**). Second, TREK1 steady-state currents, which characterize the extent of inactivation, were small in the absence of Piezo1 (5.1% (3.0 ー 10.9%, n=24)), but prominent when Piezo1 was co-expressed (77.9% (65.1 ー 88.1%, n=106); **Figure 1G**). Third, co-expression of Piezo1 delayed the time for TREK1 currents to reach their peak. Specifically, 10-90% rise times were slowed from 12.3 ms (10.5 ー 18.4 ms, n=25) to 64.6 ms (31.0 ー 95.2 ms, n=106) (**Figure 1H**). However, when we averaged all baseline-subtracted raw currents, we noticed that Piezo1 in fact accelerates the onset of TREK1 currents (**Figure S2**). We speculate that much of the delayed time-to-peak in the presence of Piezo1 results from the near lack of TREK1 inactivation: the prolonged TREK1 open times will necessarily lengthen the total rising phase of the current. We therefore chose to limit all subsequent analyses to the effects of Piezo1 on TREK1 current amplitudes and steady-state currents. As a first conclusion, our experiments fully recapitulated and confirmed the results from Glogowska and colleagues that Piezo1 has a profound modulatory effect on TREK1^27^.

As a first step to probe our hypothesis that Piezo1 modulates TREK1 through conformational signaling, we investigated the single Piezo family member from *Drosophila melanogaster* (fly Piezo), which shares only ∼30% sequence identity with mouse Piezo1^30,31^. Fly Piezo has a 5- fold smaller single-channel conductance than mouse Piezo1, and correspondingly 5-fold smaller macroscopic currents, such that the two orthologs express at similar densities (**Figure S3;** see **Methods**). We therefore reasoned that fly Piezo may provide us with initial mechanistic clues about the relative importance of structural conformation versus ion permeation. To our surprise, co-expression of fly Piezo potentiated TREK1 current amplitudes even more substantially than did mouse Piezo1. The effect was pronounced and increased with pressure, reaching an almost 15-fold potentiation at -80 mmHg (TREK1 alone: 18.3 pA (7.0 ー 38.9 pA, n=58); TREK1 + fly Piezo: 204.5 pA (57.8 ー 602.0 pA, n=85); **Figure 2A-D,F**). Fly Piezo also increased TREK1 steady-state currents to a similar extent (TREK1 alone: 5.1% (3.0 ー 10.9%, n=24); TREK1 + fly Piezo: 85.3% (66.8 ー 93.3%, n=80); **Figure 2A,E**). These data show that fly Piezo promotes TREK1 opening more strongly, and further, that mouse Piezo1 does not fully saturate the extent of TREK1 modulation. The result also supports our initial hypothesis that Piezo conformation may play a greater role than permeation in modulating TREK1, further motivating us to test their respective importance more carefully.

**Figure 2.**
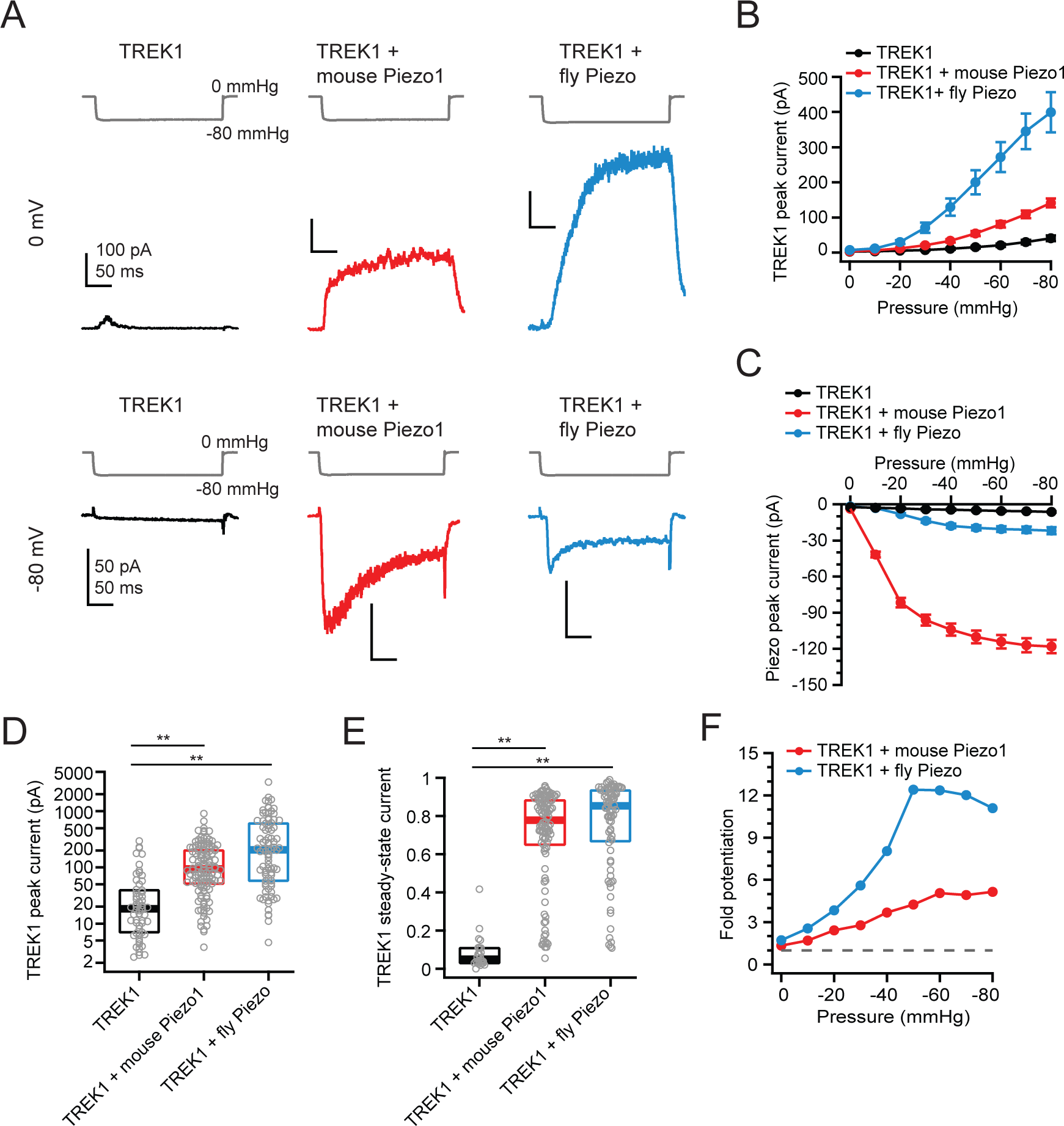
Fly Piezo potentiates TREK1 to a greater extent than mouse Piezo1. **A**, Pressure protocol (gray) and currents recorded from Neuro2A-Piezo1ko cells expressing TREK1 (black), TREK1 + mouse Piezo1 (red), or TREK1 + fly Piezo (blue). Pressure step was to -80 mmHg and voltage was 0 mV (top) or -80 mV (bottom). **B**, Mean pressure-response curves at 0 mV, generated from full pressure step protocols for TREK1 (black, n=58), TREK1 and mouse Piezo1 (red, n=121), and TREK1 + fly Piezo (blue, n=85). **C**, as in B, for inward currents at -80 mV. **D**, Box-and-whisker plots for peak outward currents elicited by a pressure step to -80 mmHg at 0 mV. **E**, Box-and-whisker plots for TREK1 steady-state current at -80 mmHg for cells expressing TREK1 (black, n=24), TREK1 + mouse Piezo1 (red, n=106), and TREK1 + fly Piezo (n=79). **F**, Fold potentiation of TREK1 peak currents by mouse Piezo1 (red) and fly Piezo (blue). Significance assessed with a Mann-Whitney U test; ∗∗p < 0.0001. Error bars are ±SEM.

### Piezo1 modulation of TREK1 signaling is independent of ion permeation

We first focused on Piezo1 ion permeation. Glogowska and colleagues already demonstrated that TREK1 modulation persists under conditions of zero net ion flow through Piezo1 and in the absence of calcium ions^27^. However, to test our hypothesis we needed to carefully quantify how the extent of modulation varied with ionic species; specifically, we focused on calcium, which regulates many cellular processes, including metabolism of phosphoinositides and other lipids, any of which could modulate TREK1 activity^32,33^.

To this end, we first used a previously described set of mutations (SNCISESEE-9K; Piezo1_9K_) that line the intracellular lateral portals of the channel and increase the relative permeability of chloride (P_Cl_/P_Na_) 50-fold, thus turning Piezo1 into a chloride-selective channel and virtually eliminating cation permeation^12^ (**Figure S4A**). Stretch-activated currents through Piezo1_9K_ at -80 mV were small compared to wild-type Piezo1 (Piezo1 alone: -115.5 pA (-75.8 ー -140.7 pA, n= 25); Piezo1_9K_ alone: -19.6 pA (-14.8 ー -28.5 pA, n=12); **Figure 3A, C-D**). Importantly, Piezo1_9K_ currents reversed near 0 mV (-7.1±1.3 mV, n=10; **Figure S4B-C**), such that TREK1 currents at this potential remain largely uncontaminated by Piezo1_9K_ currents.

**Figure 3.**
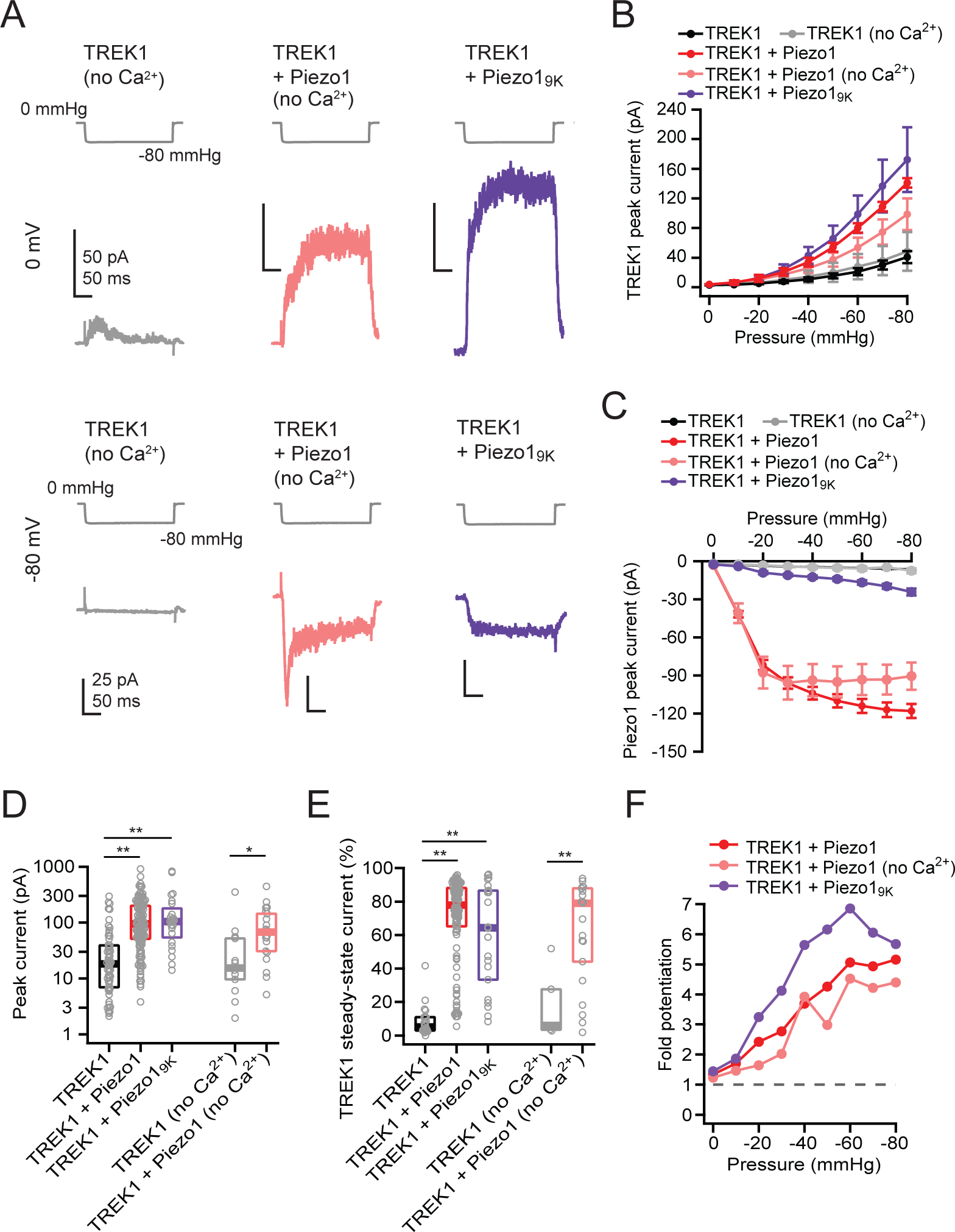
Piezo1 modulation of TREK1 is independent of ion permeation. **A**, Pressure protocol (gray) and currents recorded from Neuro2A-Piezo1ko cells expressing TREK1 (gray), or TREK1 + Piezo1 (pink) using a Ca^2+^-free pipette solution, or from cells expressing Piezo1_9K_ + TREK1 using a standard pipette solution (purple). Pressure steps were to -80 mmHg and holding potential was either 0 mV (top) or -80 mV (bottom). **B**, Mean pressure- response curves at 0 mV, generated from full pressure protocols. n=58 (TREK1), n=13 (TREK1 no calcium), n=121 (TREK1 + Piezo1), n=22 (TREK1 + Piezo1 no calcium), n=24 (TREK1 + Piezo1_9K_). **C**, as in B, for currents at -80 mV. **D**, Box-and-whisker plots for peak outward current elicited by a pressure step to -80 mmHg at 0 mV. **E,** Box-and-whisker plots for TREK1 steady- state current at -80 mmHg for cells expressing TREK1 (black, n=24), TREK1 with no calcium (gray, n=5), TREK1 + Piezo1 (red, n=106), TREK1 + Piezo1 no calcium (pink, n=20), and TREK1 + Piezo1_9K_ (purple, n=22). **F**, Fold potentiation of TREK1 peak currents by mouse Piezo1 (red), mouse Piezo1 no calcium (pink), and Piezo1_9K_ (purple). Significance assessed with a Mann-Whitney U test; ∗p < 0.001 and ∗∗p < 0.0001. Error bars are ±SEM.

Despite the reduced amplitudes of Piezo-mediated currents elicited in cells expressing Piezo1_9K_, TREK1 currents were potentiated equivalently to cells expressing wild-type Piezo1. Specifically, TREK1 peak current amplitudes were potentiated ∼7-fold (TREK1 + Piezo1_9K_: 103.5 pA (54.2 ー 185.5 pA, n=24); **Figure 3A-B,F**), and steady-state currents were increased substantially (TREK1 + Piezo1_9K_: 64.4% (33.3 ー 86.5%, n=22); **Figure 3E**). This finding further confirms that the drastically reduced permeation of cations, including calcium, does not prevent Piezo1_9K_ from modulating TREK1.

To further rule out additional potential sources of extracellular calcium that may exist downstream of Piezo activity, we also replaced 1 mM Ca^2+^ in our standard pipette solution with 1 mM Mg^2+^. In the absence of extracellular calcium, TREK1 current amplitudes were still potentiated ∼4 fold by Piezo1, from 15.3 pA (9.6 ー 52.3 pA, n=13) to 67.2 pA (31.0 ー 145.0 pA, n=22) (**Figure 3A-D,F**). Similarly, Piezo1 co-expression increased TREK1 steady-state currents from 6.1% (3.5 ー 27.7%, n=5) to 74.3% (38.5 ー 87.0%, n=19) (**Figure 3E**). We therefore conclude that the magnitude of the modulatory effect does not depend on the number and/or species of ions permeating Piezo1.

### Piezo1 conformational flexibility is specifically required for TREK1 modulation

We next focused on Piezo1 conformational changes. Structural studies have revealed two principal gating motions of Piezo channels: a flattening of the blades and a rotation of the cap^19,20,34,35^. As a means to acutely control both gating motions, we turned to two double-cysteine mutants we had previously engineered and characterized in detail (**Figure 4A**), in which mechanical gating can be controlled chemically^36^. The first construct (Piezo1_RE-CC_) allows the formation of intersubunit disulfide bonds between the blade (R1761C) and the cap (E2257C): In the absence of DTT the residues are cross-linked, which completely prevents channel gating, but in the presence of DTT this bond is reduced and the protein functions normally. Because our previous characterization of Piezo1_RE-CC_ was performed with poke stimulation, we additionally confirmed that the chemical control of Piezo1_RE-CC_ gating is maintained with pressure clamp stimulation (**Figure 4B,F**; see **Methods**). In addition, we validated that DTT did not affect either TREK1 or wild-type Piezo1 currents, or the ability of wild-type Piezo1 to potentiate TREK1 (**Figure 4D-E**).

**Figure 4.**
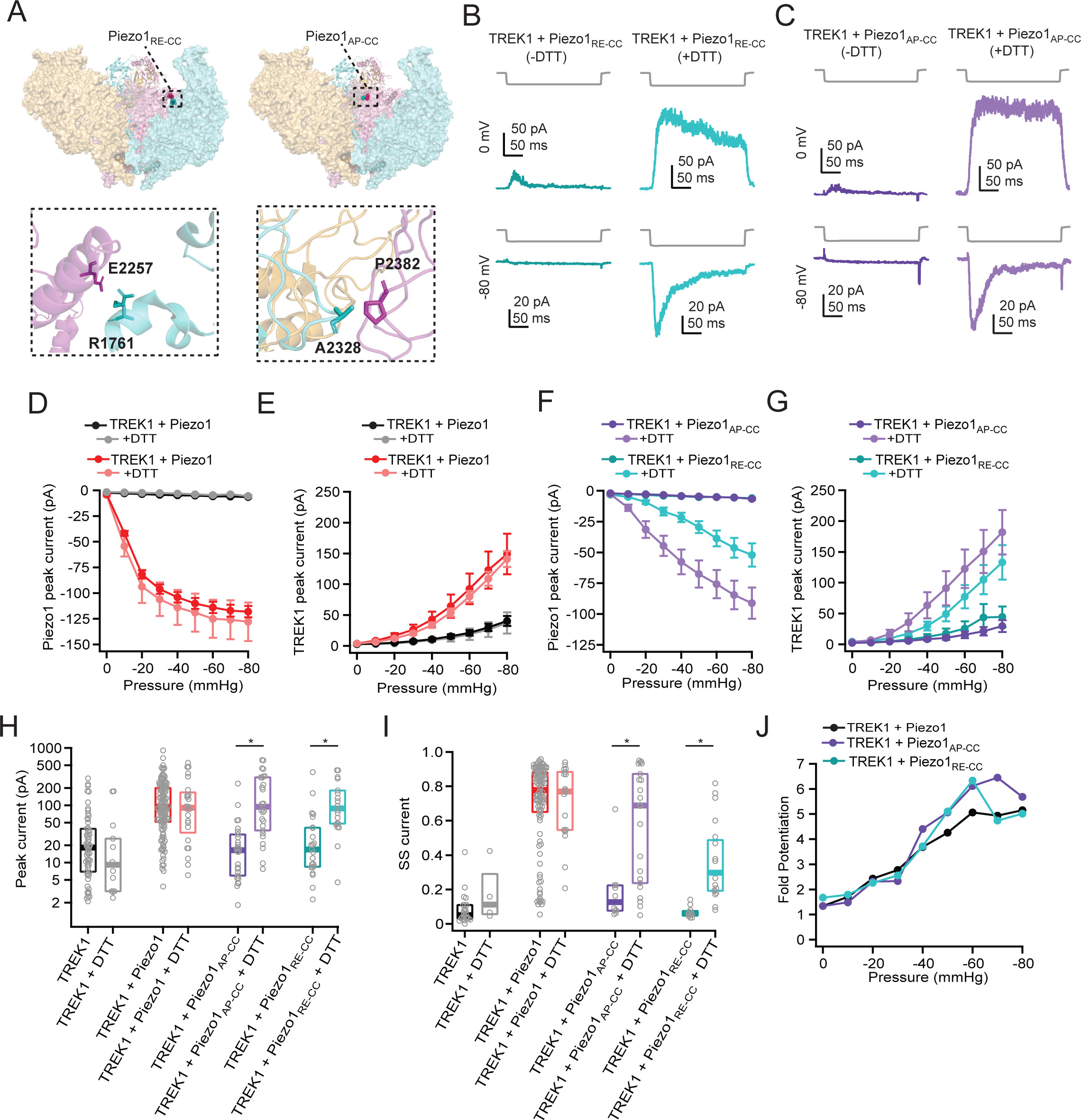
Piezo1 conformational flexibility is specifically required for TREK1 modulation. **A**, Left, top-down view of a structural model of mouse Piezo1 (PDB: 6B3R^19^) highlighting residues R1761 and E2257 at the base of the cap domain. Inset, magnified to show interacting side chains. Right, highlighting residues A2328 and P2382. **B**, Pressure protocol (gray) and currents recorded from Neuro2A-Piezo1ko cells expressing TREK1 + Piezo1_RE-CC_ in the absence (dark cyan) and presence (light cyan) of 10 mM DTT in the patch pipette. Pressure step was to -80 mmHg and voltage was 0 mV (top) or -80 mV (bottom). **C**, Same as in B for cells expressing TREK1 + Piezo1_AP-CC_. **D**, Mean pressure-response curves at -80 mV, generated from full pressure protocols, for cells expressing TREK1 (black, -DTT, n=58; gray, +DTT, n=13) and cells expressing TREK1 + Piezo1 (red, -DTT, n=121; pink, +DTT, n=24). **E**, as in D, for currents at 0 mV. **F**, As in D for cells expressing TREK1 + Piezo1_RE-CC_ (dark cyan, -DTT, n=23; light cyan, + DTT, n=21) and cells expressing TREK1 + Piezo1_AP-CC_ (dark purple, -DTT, n=26; light purple, +DTT, n=26**). G**, As in F, for currents at 0 mV. **H**, Box-and-whisker plots for peak outward currents at 0 mV and -80 mmHg. **I**, Box-and-whisker plots for TREK1 steady-state current at -80 mmHg for TREK1 (black, -DTT, n=24; gray, +DTT, n=4), TREK1 + Piezo1 (red, -DTT, n=106; pink, +DTT, n=20), TREK1 + Piezo1_RE-CC_ (dark cyan, -DTT, n=11; light cyan, +DTT, n=16) and TREK1 + Piezo1_AP-CC_ (dark purple, -DTT, n=10; light purple, +DTT, n=22). **J**, Fold potentiation of TREK1 peak currents by mouse Piezo1 (black), Piezo1_RE-CC_ (cyan) and Piezo1_AP-CC_ +DTT (purple). Significance assessed with a Mann-Whitney U test; ∗p < 0.001. Error bars are ±SEM.

We then used the chemical control of Piezo1_RE-CC_ to probe the effects of acutely restricting conformational flexibility on its ability to potentiate TREK1. In the absence of DTT, TREK1 peak currents were small (16.9 pA (8.5 ー 40.6 pA, n=23)) and inactivated nearly completely (steady-state current: 5.9% (4.8 ー 7.4%, n=11); **Figure 4B,G-I**). However, in the presence of 10 mM DTT, TREK1 peak currents were again potentiated ∼5-fold and inactivation was substantially reduced (peak current: 87.9 pA (48.0 ー 181.0 pA, n=21), steady-state current: 29.8% (19.2 ー 48.7%, n=16); **Figure 4B,G-J**). Altogether, since Piezo1_RE-CC_ gates nearly instantaneously upon application of DTT, we conclude that conformational flexibility between the cap and blades is required for modulation of TREK1.

We also tested a second construct (Piezo1_AP-CC_), which allows the formation of intersubunit disulfide bonds between two loops at the base of the cap (**Figure 4A**)^36^. When co-expressed with Piezo1_AP-CC_, TREK1 currents in the absence of DTT were small (16.5 pA (5.9ー31.0 pA; n=26)) and had small steady-state currents (12.7% (7.6 ー 22.4%; n=10)), consistent with the restricted conformational flexibility of Piezo1_AP-CC_ (**Figure 4C,G-I**). However, as expected, inclusion of DTT in the patch pipette restored potentiation of TREK1 peak currents nearly equivalently to wild-type Piezo1 (TREK1 peak current: 93.8 pA (36.5ー307 pA; n=26); steady-state current: 68.8% (23.7 ー 87.3%; n=22); **Figure 4G-J**).

Together, these experiments demonstrate that full conformational flexibility, including free movement within the cap and the flattening of the blades, is required for Piezo1 to potentiate TREK1. Further, we conclude that Piezo1 exerts its effects on TREK1 rapidly (i.e., within seconds or faster) and not through chronic effects of Piezo1 expression on cell function. Altogether, our results thus far demonstrate that Piezo1 fulfills the hallmark of conformational signaling: conformational changes alone, in the absence of ion permeation, can acutely affect other proteins.

### Piezo1 and TREK1 are not close enough to bind

What mechanism may underlie Piezo1 conformational signaling? Ca_v_1.2 calcium channels and NMDA receptors achieve conformational signaling by direct binding to downstream targets^14,16^. Further, confocal imaging and pull-down assays suggested that TREK1 and Piezo1 can physically interact when both are overexpressed at high levels^27^. We therefore hypothesized that Piezo conformational signaling may likewise require close association. To test carefully whether TREK1 channels and Piezo1 channels co-localize, a prerequisite for binding, we used stimulated emission-depletion (STED) microscopy to visualize their spatial distributions with super-resolution precision (∼80 nm; see **Methods**). We reasoned that endogenous levels of Piezo1 expression would be ideal for this query, as at low densities, in the absence of binding or other attractive force between the channels, co-localization is unlikely. To this end, we used CRISPR/Cas9 technology to insert an extracellular Myc tag into the Piezo1 gene in Neuro2A cells, which endogenously express Piezo1 at a relatively low density of ∼1-2 channels/μm^2^ (Neuro2A-Piezo1_Myc_ cells, **Figure S5A-G**)^28,37^. To visualize TREK1 channels, we inserted an extracellular HA tag into the TREK1 plasmid (TREK1_HA_; **Figure S6A-C**). In this construct, pressure-evoked peak current amplitudes were slightly smaller than for wild-type TREK1 (TREK1_HA_: 6.8 pA (4.6 ー 14.1 pA, n=26)), potentially reflecting reduced expression levels, but were potentiated by Piezo1 to a similar extent (TREK1_HA_ + Piezo1: 53.0 pA (25.7 ー 83.5 pA, n=26)), thus justifying the utility of this construct for investigating the mechanism underlying TREK1 potentiation (**Figure S6D-H**).

We then performed two-color STED microscopy on Neuro2A-Piezo1_Myc_ cells expressing TREK1_HA_. None of the images we collected showed an obvious manifestation of co-localization of TREK1_HA_ and Piezo1_Myc_ (**Figure 5A**). To quantify co-localization in an unbiased manner, we segmented STED images to obtain precise locations of individual TREK1_HA_ and Piezo1_Myc_ puncta and calculated the nearest-neighbor distance from the center of mass coordinates of each Piezo1 punctum to the nearest TREK1 punctum (NND_P1-T1_) (**Figure 5A-B; Supplemental Table 1**). To carefully probe for any hint of binding, we reasoned that if TREK1 channels were binding Piezo1 channels, the absolute maximum center of mass separation would be no larger than 30 nm (see **Methods**). Importantly, when we examined all Piezo1 puncta across three cells, only 1.2% of Piezo1 puncta had at least one TREK1 punctum within 30 nm (7 of 580 puncta; **Supplemental Table 1**). Further, when we closely inspected NND_P1-T1_ histograms, there was no hint of overrepresentation in any bin under 100 nm, which we would have expected if channels were bound (**Figure 5C)**. Thus, on average, even when TREK1 channels are in excess, the vast majority of TREK1 and Piezo1 channels are not near each other on a molecular scale; i.e., the channels do not bind at these densities.

**Figure 5.**
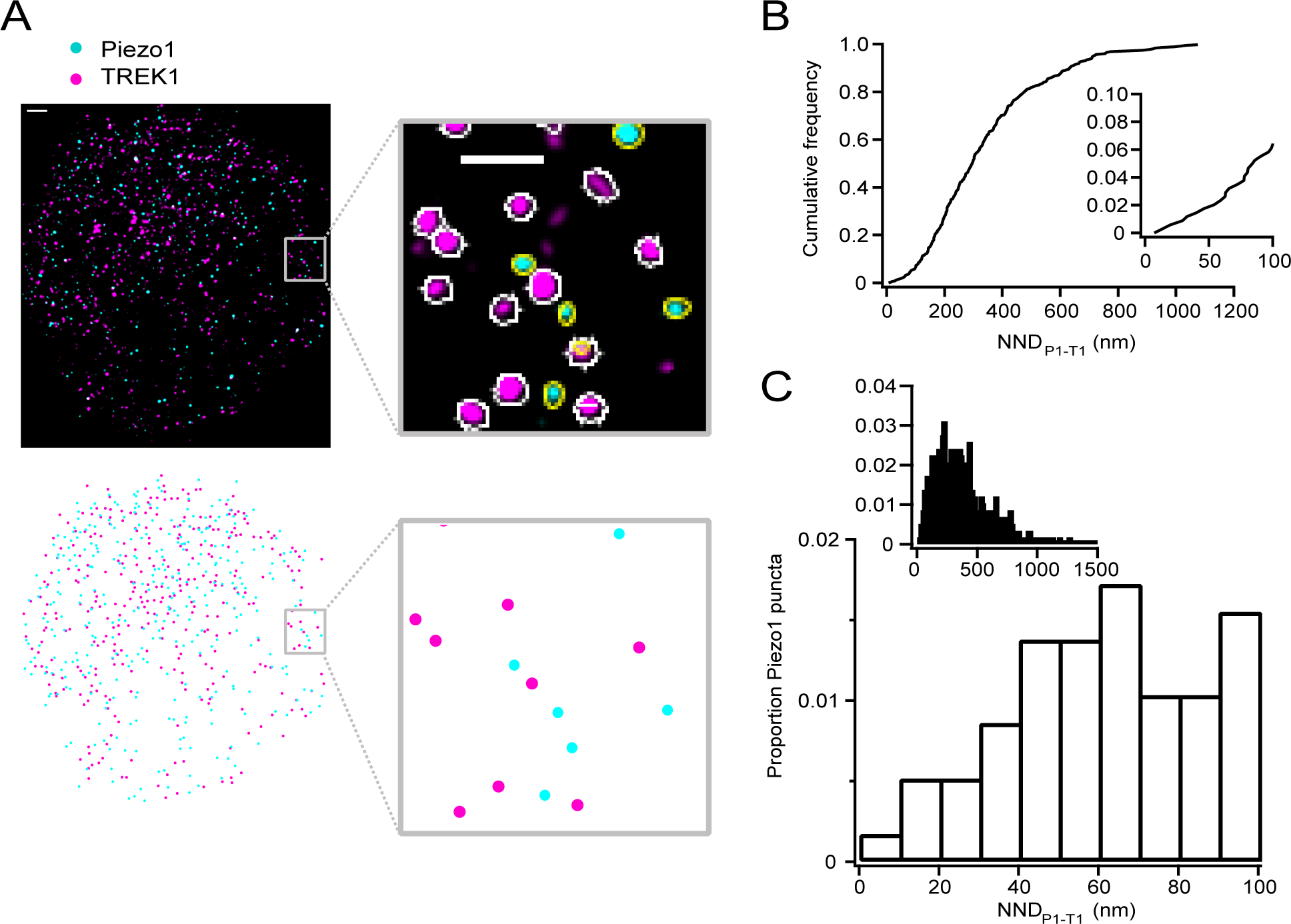
TREK1 channels do not co-localize with Piezo1 channels. **A**, Top left, Representative STED image from a Neuro2A-Piezo1_Myc_ cell (cyan puncta) overexpressing TREK1_HA_ (magenta puncta). Scale bar is 1 μm. Top right, inset showing outlines (white, yellow) of identified puncta. Scale bar is 500 nm. Bottom panels show schematized centers of mass for both Piezo1 (cyan) and TREK1 (magenta) puncta. **B**, Cumulative frequency distribution of Piezo1-TREK1 nearest neighbor distances (NND_P1-T1_) for the image shown in A (n=344 Piezo1 puncta). **C**, Histogram of NND_P1-T1_ for all images (n=3 cells, 580 puncta).

### TREK1 potentiation scales with Piezo1 density

As an alternative mechanism to direct binding, we hypothesized that a footprint-mediated mechanism could enable Piezo1 to modulate TREK1 activity over moderate distances. The estimated characteristic decay length of the Piezo footprint in a typical biological membrane is ∼14 nm, meaning the footprint may extend over a distance as far as ∼100 nm from the central pore^24,28^ (see **Methods**). If Piezo1 conformational signaling is mediated by its extended membrane footprint, then the magnitude of TREK1 potentiation should scale with increasing Piezo1 density, as more TREK1 channels will be located within a Piezo1 footprint. Indeed, we found that in wild-type Neuro2A cells, which natively express Piezo1 at low levels (1-2 channels/μm^2^)^28^, TREK1 potentiation is very modest (peak current: 114 pA vs 18 pA, p<0.001; steady-state current: 17.0% vs 5.1%, n=58,34, p<0.001; **Figure 6A-C**). To quantify more precisely how TREK1 modulation depends on Piezo1 membrane density, we next analyzed patches from Neuro2A-Piezo1ko cells in which both Piezo1 and TREK1 were overexpressed (**Figure 1**). Specifically, we calculated the number of Piezo1 and TREK1 channels in each patch from their macroscopic and single channel currents, then divided this value by the patch surface area to obtain channel densities (see **Methods**). We found that TREK1 densities and TREK1 steady-state currents both scale with Piezo1 density, showing a baseline (no substantial modulation) below ∼2 Piezo1 channels/μm^2^, half-maximal modulation at ∼5-7 Piezo1 channels/μm^2^, and saturation beyond ∼10 Piezo1 channels/μm^2^ (**Figure 6D-F)**.

**Figure 6.**
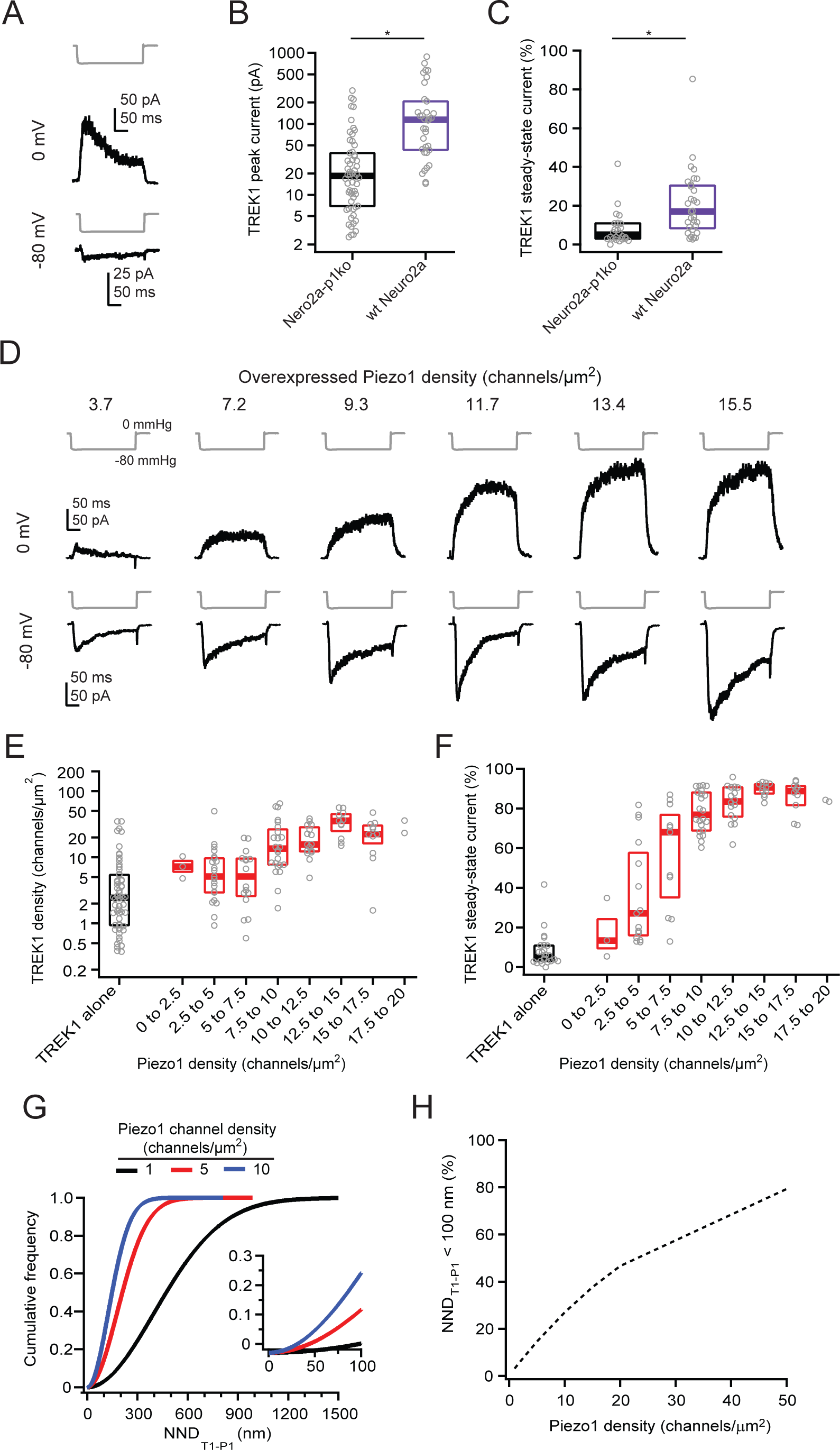
TREK1 potentiation scales with Piezo1 density. **A**, Pressure protocols (gray) and currents recorded from wild-type Neuro2A cells expressing TREK1. Pressure step was to -80 mmHg and voltage was 0 mV (top) or -80 mV (bottom). **B**, Peak TREK1 current amplitudes for wild-type Neuro2A cells (n=34) or Neuro2A-Piezo1ko cells (n=58) overexpressing TREK1. **C**, TREK1 steady-state current at -80 mmHg for wild-type Neuro2A cells (n=31) or Neuro2A-Piezo1ko cells (n=24) overexpressing TREK1. **D**, Pressure protocols (gray) and currents recorded from Neuro2A-Piezo1ko cells overexpressing TREK1 + Piezo1 at indicated Piezo1 channel densities. Pressure step was to -80 mmHg and voltage was 0 mV (top) or -80 mV (bottom). **E**, TREK1 channel density as a function of Piezo1 channel density in Neuro2A-Piezo1ko cells overexpressing both channels. **F**, TREK1 steady-state current at -80 mmHg as a function of Piezo1 channel density. **G**, Cumulative frequency distributions of nearest-neighbor distances of spatially randomly distributed TREK1 channels to randomly distributed Piezo1 channels (NND_T1-P1_) at Piezo1 channel densities of 1, 5, and 10 channels/μm^2^. **H**, Proportion of randomly distributed TREK1 channels with a Piezo1 channel within 100 nm (NND_T1-P1_ < 100 nm) as a function of Piezo1 channel density. ∗p < 0.001.

To interpret this result, we next turned to stochastic simulations to calculate the predicted nearest-neighbor distance for randomly placed TREK1 channels to their nearest Piezo1 channel (NND_T1-P1_) as a function of Piezo1 density. If TREK1 and Piezo1 are separated over >100 nm at a given density, any footprint-mediated mechanism would be implausible. At low Piezo1 densities, channels are predicted to be spatially separated: the simulations yielded that at a Piezo1 density of 1 channel/μm^2^, a TREK channel will have a median NND_T1-P1_ of 472 nm from its nearest Piezo1 channel and have only a 3% chance of residing within 100 nm of a Piezo1 channel (i.e., within its extended membrane footprint). However, at Piezo1 densities that saturate TREK1 potentiation (10 channels/μm^2^), NND_T1-P1_ decreases to 148 nm and a TREK channel has a 27% chance of residing within 100 nm of a Piezo1 channel (**Figure 6G-H**). These results suggested to us that a footprint-mediated mechanism should be investigated more thoroughly: while the fraction of TREK1 channels predicted to be within the Piezo1 footprint is relatively small, even at densities that saturate TREK1 potentiation, deviations from spatially random distributions could substantially increase this proportion.

### TREK1 channels are enriched within the Piezo1 footprint

To further address the spatial distributions of TREK1 and Piezo1, we performed additional imaging experiments, but now at Piezo1 densities that potentiate TREK1, to answer two questions: First, are TREK1 and Piezo1 channels randomly distributed with respect to each other? Second, at Piezo1 densities that modulate TREK1, is a substantial fraction of TREK1 channels close enough to feel the Piezo1 footprint?

To visualize Piezo1 channels at high densities, we used a previously characterized mouse Piezo1 construct containing an extracellular Myc tag^37^ (Piezo1_Myc_; **Figure S5H-I**). We then performed two-color STED microscopy on Neuro2A-Piezo1ko cells over-expressing TREK1_HA_ and Piezo1_Myc_ (**Figure 7A, Supplemental Video 1**). STED images were segmented to yield distributions of TREK1_HA_ and Piezo1_Myc_ puncta that corresponded to lower densities (mean±S.D.: 3.3±1.1 puncta/μm^2^ and 4.6±1.9 puncta/μm^2^, respectively; n=9 cells; **Supplemental Table 2**) than predicted from electrophysiology experiments on their respective wild-type constructs (∼10-15 channels/μm^2^), potentially resulting from reduced expression of tagged constructs, incomplete resolution of fluorescently labeled proteins at our predicted final resolution of ∼80 nm, photobleaching, and/or our stringent criteria for point segmentation (**Figure 7A**, see **Methods**). However, we noticed that in each cell, TREK1 and Piezo1 puncta densities varied locally. We therefore devised an analysis to assess the spatial distributions of TREK1 and Piezo1 locally and across all Piezo1 densities: We calculated for each TREK1 punctum not only its distance to the nearest Piezo1 punctum (NND_T1-P1_), but also the local Piezo1 density within an area of 1 μm^2^ (**Figure 7B**, see **Methods**). The data showed that NND_T1-P1_ decreased as a function of local Piezo1 density for all cells, from ∼600-800 nm at 1 punctum/μm^2^ to ∼100 nm at 10 puncta/μm^2^ (**Figure 7C-D**, n=9 cells). Further, individual distributions for each cell overlapped, and we therefore pooled puncta from all cells for subsequent analyses.

**Figure 7.**
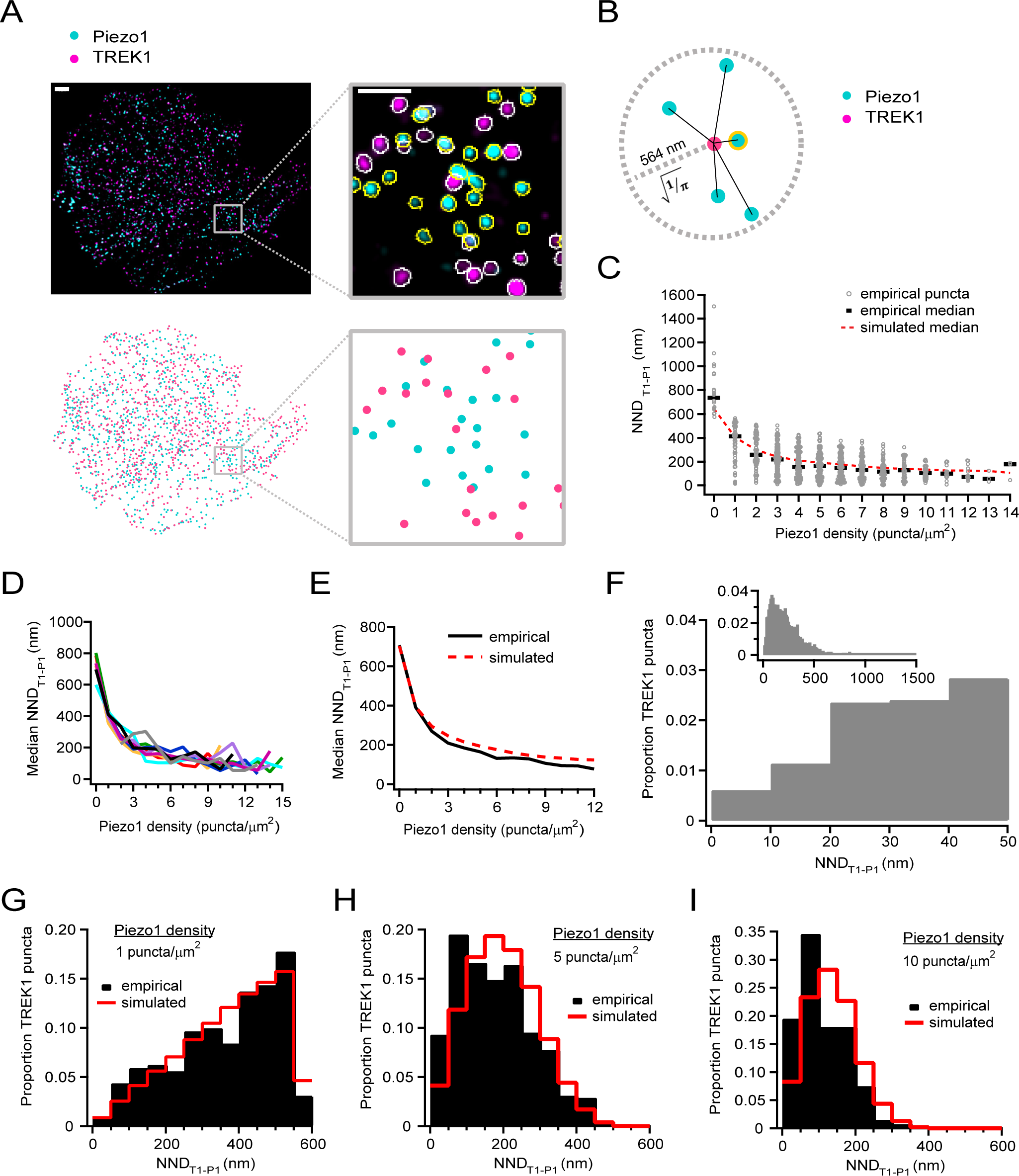
TREK1 channels are enriched within the predicted Piezo1 footprint. **A**, Top left, Representative STED image from a Neuro2A-Piezo1ko cell overexpressing TREK1_HA_ (magenta) + Piezo1_Myc_(cyan). Scale bar is 1 μm. Top right, inset showing outlines (white, yellow) of identified puncta. Scale bar is 500 nm. Bottom panels show schematized centers of mass for both Piezo1 (cyan) and TREK1 (magenta) puncta. **B**, Schematic illustrating calculation of local Piezo density (count of cyan puncta) and nearest neighbor (NND_T1-P1_, yellow highlighted punctum) for a central TREK1 punctum (magenta) within a radius of 564 nm (1/√π). **C**, NND_T1-P1_ as a function of local Piezo density for the image shown in A. N=895 puncta. Red dashed line represents median of 1,000 simulations of randomly distributed TREK1 channels (n=894,158 puncta). **D**, Median NND_T1-P1_ as a function of local Piezo1 density for 9 cells. **E**, Median NND_T1-P1_ as a function of local Piezo1 density for all empirical puncta (n=3,950 puncta, N=9 cells) and simulated puncta (n=3,944,121 puncta). **F**, Histogram of NND_T1-P1_ for all puncta (n=3,950 puncta). **G-I**, As in F, for TREK1 puncta with a local Piezo1 density of 1, 5 and 10 Piezo puncta/μm^2^. N=321, 451, and 133 puncta, respectively.

To first ask if TREK1 channels and Piezo1 channels are randomly distributed, we compared NND_T1-P1_ distributions to more sophisticated stochastic simulations, in which we maintained the respective shape (outline) of each cell and its exact Piezo1 locations and randomized all TREK1 locations, while keeping their densities unaltered (**Figure 7E**; see **Methods**). Surprisingly, this analysis revealed that the empirical NND_T1-P1_ was systematically lower than predicted stochastically. Specifically, at low Piezo1 densities (1 punctum/μm^2^) the effect is negligible (median NND_T1-P1_ 391 nm vs 394 nm; n=321 puncta), but at medium Piezo1 densities (5 puncta/μm^2^) it is pronounced (median NND_T1-P1_ 165 nm vs 194 nm; n=451 puncta), and at high Piezo1 densities (10 puncta/μm^2^) it is striking (median NND_T1-P1_ 95 nm vs 132 nm; n=133 puncta) (**Figure 7G-I**). Thus, our analysis revealed a spatial bias between TREK1 and Piezo1: at Piezo1 densities where conformational signaling is saturating, TREK1 and Piezo1 are substantially closer than predicted by randomness. Still, despite this spatial bias, even at Piezo1 densities that fully saturate TREK1 potentiation (10 puncta/μm^2^), fewer than 5% (162/3,945 puncta; **Figure 7F**) of TREK1 puncta had their nearest Piezo1 punctum within 30 nm, and likewise, the fraction of Piezo1 puncta with a TREK1 punctum within 30 nm was similarly low (7.5%, 162/4,897 puncta). These results confirm our above conclusion that TREK1 and Piezo1 channels do not co-localize (i.e., bind).

Finally, with the goal of assessing the plausibility of a footprint-mediated mechanism for conformational signaling, we asked if a substantial fraction of TREK1 channels is located within the Piezo1 footprint. We found that at low Piezo1 density (1 punctum/μm^2^), where conformational signaling is absent, only ∼5% of TREK1 puncta had a Piezo1 punctum within 100 nm - the maximal range we reasoned for the Piezo1 membrane footprint (**Figure 7G**). At an intermediate Piezo1 density (5 puncta/μm^2^), where conformational signaling is half-maximal, this value rose to nearly 30% (**Figure 7H**). Most strikingly, at Piezo1 densities that begin to saturate conformational signaling (10 puncta/μm^2^), more than 50% of TREK1 puncta had a Piezo1 punctum within 100 nm (**Figure 7I**). This quantification demonstrates that at Piezo1 densities that matter functionally for conformational signaling, most TREK1 channels reside within a predicted Piezo1 footprint. We therefore propose that a footprint-mediated mechanism is plausible and must be considered when further investigating Piezo conformational signaling.

## Discussion

In this study, we aimed to answer if Piezo1 channels are capable of conformational signaling. By using as a readout the modulation of K2P channels by Piezos, first identified by the Honoré group^27^, we found that full conformational flexibility of Piezo1, but not ion permeation through it, is required for its modulation of TREK1. This finding conceptually expands the mechanisms by which mechanical forces are transduced into cellular signals.

Several limitations of our experimental approaches need to be considered: First, we do not know exactly how chemical crosslinking biases the conformation of Piezo1. However, an all-atom molecular dynamics simulation of Piezo1 found that contact and separation of A2328- P2382 and R1761-E2257 occur during the tension-induced gating transition. This is fully supported by the comparison of partially flattened and curved cryo-EM structures^34,38^. Thus, it is reasonable to predict that chemical crosslinking inhibits both blade flattening and cap rotation. Nanoscopic fluorescence imaging of Piezo1 cysteine mutants^23^, or measurements of the shape of lipid bilayer vesicles containing them^26^, which have both been performed on wild-type Piezo1, could further probe the structures of both mutants in the presence and absence of DTT. Second, the spatial resolution of our STED imaging approach is limited to ∼80 nm. This leads to an underestimation of channel density for channels labelled with the same fluorophore; however, the positional accuracy for locating TREK1 and Piezo1 is higher, as they are labeled with two different fluorophores. Third, it is possible that TREK1 and Piezo1 spatial distributions are dynamic; indeed, Piezo1 channels diffuse rapidly in the membrane^39^. Moreover, flattening of Piezo1 and its membrane footprint may lead to a rapid attraction or repulsion of TREK1, and thus a change in their relative distributions upon Piezo1 opening that would not be captured by our static imaging paradigm. Fourth, our estimate of the membrane footprint radius (100 nm) is derived from the most curved conformation of Piezo1 obtained in a cell-free system^24^. In a biological membrane, the Piezo1 resting state likely adopts a more expanded conformation, resulting in a larger protein radius, albeit a less pronounced membrane curvature and correspondingly smaller footprint, giving rise to error in our estimate. In addition, the membrane bending modulus and presence of other membrane-associated proteins vary locally, and therefore the precise geometry of the footprint differs for each individual Piezo channel^25,26^. Finally, our observation that TREK1 and Piezo1 are statistically closer than predicted by randomness may arise from non-specific mechanisms (e.g., biased membrane protein localization in the membrane through lipid rafts), Piezo1-specific biological mechanisms (e.g., Piezo1 channels attracting TREK1 channels), or from technical limitations (e.g. membrane ruffling), and we currently do not know which mechanism is at play.

How does Piezo1 conformational signaling complement ionotropic signaling? First, Piezo1 conformational signaling is rapid. The chemical control of Piezo1 gating we employed circumvents the potentially confounding effects of basal Piezo1 activity (e.g., via its chronic effects on cholesterol and cytoskeletal dynamics^40–42^) and directly shows that conformational signaling occurs within no more than 60 seconds - roughly the time it takes to establish and record from a cell-attached patch. Given that Piezo1 activates within milliseconds, it is entirely possible that conformational signaling is equally rapid. However, testing the detailed kinetics will require more sophisticated experiments. Second, Piezo1 conformational signaling apparently functions without direct protein-protein interactions (binding), which we would have detected as channel co-localization. Instead, our imaging data predict that conformational signaling occurs over relatively long distances: we calculated that even at Piezo1 densities that saturate TREK1 potentiation (10 channels/μm^2^), many TREK1 channels reside 50-100 nm away from their nearest Piezo1. This length scale rivals the size of calcium nanodomains (∼50 nm) known to exist around ion channel pores due to the rapid ion diffusion and natural calcium buffering capacity of the cell^43,44^. Thus, Piezo1 conformational signaling has the capability to extend spatially as far as, and potentially further than, ionotropic signaling. Third, since Piezo1 is activated (gated) at very low membrane tension, Piezo1 conformational signaling must also be considered a highly sensitive mechanism. Our data showing the fold-potentiation of TREK1 saturates near -60 mmHg directly supports this interpretation.

Our spatial analyses rule out direct binding as a mechanism for Piezo1 conformational signaling to TREK1, and further reveal that TREK1 channels are moderately enriched near Piezo1 channels, such that at Piezo1 densities that potentiate TREK1, more than half of TREK1 channels have a Piezo1 channel within 100 nm, i.e., within the Piezo1 footprint. Interestingly, our finding that fly Piezo potentiates TREK1 more strongly than does mouse Piezo1 may provide further evidence for a footprint-mediated mechanism: its relatively low homology with the mouse isoform may endow it with a different intrinsic curvature than mouse Piezo1 and consequently a different footprint; this will require future structural studies. If the footprint does mediate conformational signaling, how might it do so? We recognize at least three potential mechanisms. First, the curved Piezo footprint could bias the local concentration of lipids. Indeed, Glogowska and colleagues previously suggested a role for cholesterol reorganization, based on coarse-grained molecular dynamics simulations predicting that Piezo1 alters the local lipid environment and the known effects of cholesterol on membrane tension and K2P channel function^27,45–48^. However, as both Piezo1 and TREK1 mechanical activity are strongly modulated by cholesterol, potentially through both direct binding to the channels as well as the known effects of cholesterol on membrane curvature and tension, we find it difficult to interpret any experiments testing the ability of Piezo1 to potentiate TREK1 in varying cholesterol concentrations^27,49^. Moreover, given that Piezo1 channels, even at overexpression densities, occupy <5% of the cell membrane^28^, and cholesterol comprises up to 40%^50^, it seems unlikely that Piezo1 could substantially alter free cholesterol concentrations. Second, the curved Piezo1 footprint could induce an asymmetric bilayer pressure profile, which K2P channels are known to respond to by transitioning from a closed “down” conformation to an open “up” conformation^51–54^. Several voltage-gated ion channels are also known to be modulated by membrane curvature through their voltage-sensing domain, which makes them potential targets for Piezo1 conformational signaling^55,56^. Third, the footprint may mediate conformational signaling via the cytoskeleton that is associated with the Piezo protein and/or its footprint, and thereby modulate other force-sensitive proteins^57^. Here again, however, cytoskeletal disrupting agents affect both direct force transmission to the channels, as well as indirectly alter membrane tension and its propagation, and therefore their effects are difficult to interpret^58–60^.

Of course, these three mechanisms are not mutually exclusive and importantly, lipid and cytoskeletal modulation could both occur independently of the Piezo footprint. Alternatively, Piezo1 could influence TREK1 through a second messenger system. For example, phospholipase D2 (PLD2) rapidly (<1 ms) translocates in response to membrane stress, subsequently generating phosphatidic acid (PA) that could directly activate TREK1, though it has yet to be explored what, if any, effects Piezo1 may have on PLD2 activity^61–63^.

In our opinion, definitive answers for a complete mechanism can only be provided by a bottom-up approach: purifying, reconstituting, and functionally investigating the interaction between both proteins in an artificial bilayer system would answer what roles membrane lipids (cholesterol), second messenger systems (PLD2/PA), or indirect force transmission (cytoskeleton) play. Circumstantial evidence might also come from a candidate approach. Screening a library of membrane proteins known to be specifically sensitive to cholesterol, other lipid signaling molecules and enzymes, membrane curvature, or mechanical forces, may hint at a common underlying mechanism. Our future efforts will pursue these approaches.

What is the prevalence of Piezo conformational signaling? Based on the previous finding that mouse Piezo2 also has the ability to potentiate TREK1, as well as our discovery that the distantly related fly Piezo channel can do so to an even greater extent, it is highly probable that conformational signaling generalizes to all Piezo proteins^27^. Intriguingly, Piezos may also couple to many different signaling partners. Indeed, K2P family members TREK2 and TRAAK are also modulated by Piezo1^27^. Clearly, the full range of membrane proteins that are targets of Piezo conformational signaling and thus function as novel molecular substrates of mechanotransduction has yet to be explored. We further anticipate that some physiological functions that have previously been attributed to its ionotropic function as a channel based solely on genetic knockout of the protein could, in fact, involve Piezo conformational signaling.

## Author Contributions

A.H.L and M.E.C designed and performed experiments and analyzed data. A.H.L and J.G. wrote the manuscript.

## Acknowledgments

We thank Michael Sindoni, Will Sharp, Mona Ghimire, Jason Wu, and Michael Young for careful reading of the manuscript. Funding was provided by the Holland-Trice Scholar Program, the National Science Foundation Graduate Research Program Award, and NIH 5R01NS110552. We thank the Duke Light Microscopy Core Facility for support and the Duke Functional Genomics Core for generating Neuro2A-Piezo1_Myc_ cells. We thank Dr. Anne West and lab members and the Duke Viral Vector Core for Western Blot support.

## STAR Methods

## RESOURCE AVAILABILITY

### Lead contact

Further information and requests for resources and reagents should be directed to and will be fulfilled by the lead contact Jörg Grandl.

### Materials availability

New plasmids and cell lines generated in this study are available from the lead contact upon request.

### Data and code availability

- All data reported in this paper will be shared by the lead contact.
- All original code used in this study is available in our Github repository (https://github.com/GrandlLab?tab=repositories).
- Any additional information required to reanalyze the data reported in this paper is available from the lead contact upon request.

## EXPERIMENTAL MODEL AND SUBJECT DETAILS

### Cell lines

All cells were cultured at 37°C and 5% CO_2_ in Minimum Essential Medium (ThermoFisher Scientific) supplemented with 0.1 mM non-essential amino acids, 1 mM pyruvate, 10% FBS (Clontech), 50 U/mL penicillin, and 50 mg/mL streptomycin (Life Technologies). All cells were STR authenticated and verified negative for mycoplasma by American Type Culture Collection (ATCC).

Electrophysiology experiments were performed using Neuro2A cells CRISPR-engineered to lack endogenous expression of Piezo1, which have previously been described (Neuro2A- Piezo1ko; a gift of Gary Lewin^64^) or wild-type Neuro2A cells (ATCC#CCL-131), as indicated. Imaging experiments were performed using either Neuro2A-Piezo1ko cells or a cell line in which a Myc tag (EQKLISEEDL) was inserted via CRISPR/Cas9 technology into the Piezo1 gene of wild-type Neuro2A cells (ATCC #CCL-131) immediately following residue 2422 (Neuro2A- Piezo1_Myc_). Briefly, 100,000 Neuro2A cells were electroporated with 1 mg TrueCut Cas9 protein (ThermoFisher Scientific), 22.5 pmol sgRNA (Synthego) and 40 pmol single stranded oligonucleotide donor (Integrated Data Technologies) using the Neon system (ThermoFisher Scientific). The oligonucleotide donor was designed to incorporate the Myc epitope tag as well as two additional silent mutations to prevent re-cleavage by Cas9 after homology-directed repair (**Figure S5A-B**). After confirming the presence of successful knock-in (KI) of the Myc sequence, cells were plated at limiting dilution in 96-well plates to isolate clones. Clones were expanded for several weeks and screened by PCR using a Myc KI-specific forward primer. Positive clones were then subjected to PCR sequencing to confirm the presence of the Myc tag in the appropriate genomic location. Neuro2A-Piezo1_Myc_ cells had Piezo1-mediated stretch-induced currents that were comparable to wild-type Neuro2A cells (∼1 channel/μm^2^; **Figure S5D-E**). Piezo1 channel densities measured with STED imaging were similar to those measured with electrophysiology and consistent with our previous work, Piezo1 channels were spatially randomly distributed in this cell line (**Figure S5J-K; Supplemental Table 1**)^28^.

## METHOD DETAILS

### Cell culture and plasmids

TREK1_HA_ was generated by inserting an HA tag (YPYDVPDYA) immediately following residue N134 in the loop connecting the cap and P1 helix using the Q5 mutagenesis kit (New England Biosciences) following the manufacturer’s protocol for large insertions using non-overlapping primers and was sequence-verified (Genewiz). Mouse Piezo1_Myc_ has a Myc tag (EQKLISEEDL) immediately following residue 2422, and displays normal mechanosensitivity, as previously described^37^. To measure membrane expression levels of Piezo constructs, a Myc tag was inserted into the same location of Piezo1_RE-CC_, Piezo1_AP-CC_, and Piezo1_9K_; additionally, a stop codon was inserted at the end of the Piezo1_9K_ coding sequence to stop expression of the mRuby fusion protein and avoid spectral overall with the secondary antibody. Fly Piezo_Myc2325_ was generated by inserting a Myc tag immediately following residue N2325 and had normal mechanosensitivity (**Figure S8A-C**). Details and sources for all other plasmids are in the Key Resources Table.

For electrophysiology, Neuro2A-Piezo1ko cells were transiently transfected 40-48 hours prior to recording in 6-well plates using Lipofectamine 2000 (Thermo Fisher Scientific) according to the manufacturer’s protocol. A total of 4 μg of plasmid DNA was used for each transfection: 4 μg TREK1 alone, 4 μg Piezo construct alone, or 0.75 μg TREK1 + 3.25 μg Piezo construct. Transfected cells were reseeded onto poly-L-lysine- and laminin-coated coverslips 16-24 hours before recording. For imaging, 48 hours before immunocytochemistry, Neuro2A-Piezo1_Myc_ or Neuro2A-Piezo1ko cells were plated in 24-well plates on poly-L-lysine- and laminin-coated No. 1.5 coverslips (Warner Instruments, 64-0732) and transfected with Lipofectamine 2000 according to the manufacturer’s protocol with either 200 ng TREK1_HA_ or 200 ng GFP (Neuro2A- Piezo1_Myc_) or 150 ng TREK1_HA_ and 650 ng Piezo1_Myc_.

### Electrophysiology

Electrophysiological recordings were performed at room temperature using an EPC10 amplifier and Patchmaster software (HEKA Elektronik). Data were sampled at 10 kHz and filtered at 2.9 kHz for all experiments except measurements of single-channel conductance of TREK1 and fly Piezo, which were sampled at 50 kHz and 100 kHz respectively, filtered at 2.9 kHz during recording, then digitally filtered offline at 1 kHz prior to analysis. The cell-attached bath solution used to zero the membrane potential was (in mM): 155 KCl, 3 MgCl_2_, 5 EGTA, 10 HEPES, pH 7.35 with NaOH. Borosilicate glass pipettes (1.5 OD, 0.85 ID, Sutter Instrument Company) were filled with pipette buffer solution (in mM): 150 NaCl, 5 KCl, 2 CaCl_2_, 1 MgCl_2_, 10 HEPES, pH 7.3 with NaOH and had resistances ranging between 1.3-3 MΩ (mean±S.D. = 2.1±0.3 MΩ). In these buffers, assuming an intracellular K^+^ concentration of 120 mM, the predicted reversal potential for K^+^ is -81.6 mV. The measured reversal potential for TREK1, generated from patches with at least 100 pA of outward current at +60 mV to avoid leak effects, was -60.3±4.8 mV (n=8, **Figure S1B-C**); however, inward currents through TREK1 channels at -80 mV were negligible (5-10 pA, **Figure 1**). Negative pressure was applied through the patch pipette with an amplifier-controlled high-speed pressure clamp system (HSPC-1; ALA Scientific Instruments). Mechanical sensitivity was probed by applying brief (250 ms) pressure steps from 0 to -80 mmHg (Δ = -10 mmHg). All pressure steps were separated by 10 s to allow for recovery from inactivation. Voltage during the pressure steps was alternated between -80 mV and 0 mV at each pressure level.

For experiments involving breaking of double-cysteine bonds, 10 mM dithiothreitol (DTT) was included in the pipette solution. DTT was kept on ice and made fresh from frozen stock (1 M) hourly. The chemical control of double-cysteine mutants we previously observed using poke stimulation was maintained using pressure-clamp stimulation: In cells transfected with Piezo1_RE-CC_ or Piezo1_AP-CC_, stretch-activated currents at -80 mV were not larger than in cells transfected with TREK1 alone (TREK1 alone: -5.9 pA (-5.2 ー -7.1 pA, n=58); TREK1 + Piezo1_AP-CC_: -5.9 pA (-4.7 ー -7.3 pA, n=23); TREK1 + Piezo1_RE-CC_: -6.0 pA (-5.4 ー -8.4 pA, n=26); **Figure 4B-C**), which is consistent with the newly introduced disulfide bond preventing Piezo channels from gating. However, inclusion of 10 mM DTT in the patch pipette resulted in large stretch-activated currents through both Piezo1_RE-CC_ and Piezo1_AP-CC_ (TREK1 + Piezo1_RE-CC_ + DTT: -41.3 pA (-23.2 ー -63.3 pA, n=21); TREK1 + Piezo1_AP-CC_: 82.3 pA (-45.0 ー -125.0 pA; n=26; **Figure 4G**).

### Microscopy sample preparation

Cells were transiently transfected with TREK1_HA_ and/or Piezo1_Myc_ plasmids and plated on No 1.5 coverslips (Warner Instruments: CS-12R15, Catalog # 64-0712) 48 hours before staining, fixed in 2% formaldehyde, blocked with 10% normal goat serum, and stained with 1:100 chicken anti-Myc (Novus) primary antibody followed by 1:500 Alexa Fluor plus 594 goat anti-chicken (Thermo Fisher Scientific) secondary antibody to label Piezo1_Myc_ channels and 1:500 rabbit anti-HA (Cell Signaling Technology) primary antibody followed by 1:200 Atto 647N goat anti-rabbit (Rockland Instruments) secondary antibody to label TREK1_HA_ channels. Coverslips were then mounted with ProLong Glass Antifade Mountant (ThermoFisher Scientific) and cured at room temperature overnight before image acquisition. STED images were collected from at least three coverslips per condition, with n=3 independent transfections to generate biological replicates.

### STED image acquisition and deconvolution

Two-color STED was performed on co-labeled images of either Neuro2A-Piezo1ko cells overexpressing Piezo1_Myc_ or Neuro2A-Piezo1_Myc_ cells, both transiently transfected with TREK1_HA_. All image collection was performed on a Leica SP8 instrument equipped with a 100x/1.4 HCX PL APO OIL WD 90 μm objective, pulsed White Light Laser, and HyD detectors, using Leica Application Suite Software (3.5). The 594 red channel, corresponding to Piezo1_Myc_, was excited using 5% laser power at 591 nm, and emitted light was collected between 603 and 641 nm with 22% gain. The 647 far red channel, corresponding to TREK1_HA_, was excited using 3% laser power at 641 nm and emitted light was collected between 651 and 779 nm with 42% gain. The 647 channel was collected using 2x frame averaging, to reduce noise. A pulsed 775 nm STED depletion laser at 20% laser power in the 647 channel, with gating (0.7 to 4.2), and 70% laser power in 594 channel, with gating (0.7 to 4.2), was used to improve image resolution to ∼80 nm (**Supplemental Table 1-2**). For STED image collection, Z-stacks were acquired from the middle to the top of the cell in steps of 220 nm with a Märzhäuser linearly encoded piezo Z stage (**Supplemental Movie 1**).

All channels from the STED images were deconvolved using Huygen’s Professional (Scientific Volume Imaging). The refractive index was corrected to match the immersion oil (1.5) and images were cropped as necessary to isolate single cells. Deconvolution was then performed with an automatically generated theoretical point spread function and the preset Classic Maximum Likelihood Estimation CMLE deconvolution algorithm, with the signal-to-noise ratio set to 5.0.

### Confocal microscopy image acquisition

Confocal imaging was used to assess antibody specificity (**Figure S5F-I, Figure S6B-C**) and quantify membrane expression levels of all Myc-tagged Piezo constructs (**Figure S8D-E**). Antibody specificity was assessed using widefield confocal images at 100X magnification acquired with the same objective, excitation and emission collection parameters as STED images, as described above, but without depletion. For membrane expression quantification, images were acquired using an HC PL APO CS2 40x/1.30 Oil objective on the above described Leica SP8 scope. The 594 red channel, corresponding to Piezo1_Myc_, was collected using 5% laser power at 591 nm, and emitted light was collected between 603 and 641 nm with 13% gain. The 488 green channel, corresponding to GFP, was collected using 7% laser power at 490 nm, and emitted light was collected between 501 and 561 nm with 52% gain. Single Z-slices of the central plane, the midpoint of most cells in a field view, were collected over ∼12 images, yielding approximately 200 cells per condition.

### Western Blot

The membrane fraction was isolated from Neuro2A cells using the ProteoExtract® Subcellular Proteome Extraction Kit (Sigma) and concentrated to 1-2 mg/mL using a Pierce^TM^ concentrator (3K molecular weight cutoff; ThermoFisher Scientific). 50 μg of protein was loaded in each well of a 4-20% precast polyacrylamide gel (BioRad). Gels were transferred (1.5 A for 15 minutes) using the Trans-blot Turbo semi-dry transfer system (BioRad) onto a 0.2 μm PVDF membrane. To visualize Piezo1_Myc_, blots were stained with 1:750 chicken anti-myc (Novus) primary antibody followed by 1:2,000 Goat Anti-Chicken horseradish peroxidase (Novex) secondary antibody and visualized via enhanced chemiluminescence (ECL) with SuperSignal™ West Femto Maximum Sensitivity Substrate (ThermoFisher Scientific). To visualize the transferrin loading control, blots were stripped for 10 minutes with Restore Western Blot Stripping Buffer (ThermoFisher Scientific) and then stained again with 1:1,000 mouse anti-transferrin primary antibody (Invitrogen) followed by 1:2,000 goat anti-mouse peroxidase (Invitrogen) and again visualized using ECL.

## QUANTIFICATION AND STATISTICAL ANALYSIS

### Electrophysiology

All data were analyzed and final plots were generated using Igor Pro 8.02 (Wavemetrics). Only patches retaining a gigaohm seal after all pressure steps were included in the analysis. Peak currents were measured following baseline subtraction of the mean current 50-200 ms immediately prior to the pressure stimulus. Only patches with TREK1 peak currents >20 pA were included in the steady-state analysis and in the calculation of 10-90% rise times. TREK1 steady-state currents were measured as the mean current during the last 50 ms of the pressure pulse. All 10-90% rise times were calculated as the time for the baseline-subtracted current to rise from 10% to 90% of its peak value. To account for current noise, all 40% decay times were calculated as the time at which five consecutive current amplitude points (corresponding to 0.5 ms) were below 40% of the peak value. Fold potentiation of TREK1 currents was calculated at each pressure as the ratio of median currents ((TREK1 + Piezo)/TREK1).

Single channel current amplitudes were calculated by generating all-points histograms with binning determined using the Freedman-Diaconis method and an optimal bin width of 2*IQR(x)/N^1/3^, where IQR is the interquartile distance, N is the number of observations, and the bins are evenly distributed between the minimum and maximum values. For mouse Piezo1, we took advantage of the low open probability of Piezo1 in the absence of membrane tension to measure the single-channel current (n=3-5 openings per patch) of Piezo1 in most patches (106/121 patches; **Figure S7A-B**). Because we did not observe TREK1 single-channel openings in most patches, we used an altered protocol to measure the single-channel current of TREK1 at 0 mV in our solutions. TREK1 openings in a cell-attached configuration are brief, and therefore 7-10 baseline-subtracted openings from one patch were pooled. For both Piezo1 and TREK1, binned data were fit with a double-Gaussian equation of the form

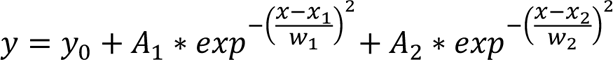

, where y_0_ is the baseline current, A_1_ and A_2_ are the peak amplitudes, x_1_ and x_2_ are the centers of the fits, and w_1_ and w_2_ their respective widths. The difference between x_1_ and x_2_ reflects the difference between mean current in the open and closed state and was used to calculate single channel currents. Because fly Piezo has a small single-channel conductance, we could not resolve openings at -80 mV ^31^. We therefore measured slope conductances for both mouse Piezo1 and fly Piezo by fitting a line to single channel currents as a function of voltage between -160 mV and -240 mV (**Figure S3**). To compare macroscopic current amplitudes and thus compare channel density between fly Piezo and mouse Piezo1, we used day-matched controls to limit variability due to passage number.

Patch dome areas were estimated from pipette resistances using a linear fit to data previously recorded in the lab with Differential Interference Contrast imaging^1,28^ (**Figure S7E**). Specifically, a plot of surface area as a function of pipette resistance was fit with a linear equation, y = 13.33 – 2.55x, which was then used to estimate patch dome surface area (y, μm^2^) for a given pipette resistance (x, MΩ). We calculated surface area of each patch from its pipette resistance, a relationship we had previously calibrated via imaging of the patch dome area^1^ (**Figure S7E**) and divided the number of channels in each patch by this value to calculate channel densities.

We found that values for peak currents, steady-state currents, and 10-90 rise times were widely distributed, sometimes over more than two orders of magnitude, and that the distributions were visibly not symmetrical. For this reason, we chose to report median values and 1st and 3rd quartiles. For box plots, boxes represent median and 1st and 3rd quartiles. All other data are reported as mean±SD or mean±SEM, as indicated.

### Image processing

Images were processed in FIJI (version 2)^65^ as 16-bit TIFF files. A single z-slice for spatial analysis from the cell surface (top) was manually chosen from the Z-stack. Noise thresholds were set for each cell by identifying the intersection of Gaussian distributions fit to the intensity histograms of segmented puncta from the surface z-slice and a separate z-slice chosen from the unlabeled interior of the cell. Manually drawn ROIs were generated around the xy perimeter of each cell, and signal outside the ROI was cleared. Both 594 (Piezo1_Myc_) and 647 (TREK1_HA_) channels were auto-enhanced and filtered with a 2.0 pixel Gaussian Blur Filter. Individual puncta from both overexpressed TREK1 and Piezo1 channel conditions were segmented using StarDist’s Versatile (fluorescent nuclei) Model with the following settings (Normalized Image Percentile 3-100, Probability/Score Threshold: 0.5, Overlap Threshold: 0.2, Number of Tiles: 1, Boundary Exclusion: 2)^66^. Automated segmentation parameters were developed based on manual segmentation. Endogenous Piezo1 puncta from Neuro2a- Piezo1_Myc_ images were manually segmented. We note that the size and intensity of individual segmented puncta in images varies, particularly for TREK1. This variance primarily stems from the variable z position along the axial point spread function of a punctum, especially given that our image acquisition settings were optimized exclusively to improve lateral resolution in x and y. Additional variance may come from two or more channels located closer to each other than the resolution limit of our experiments. The center of mass for each Piezo or TREK punctum was identified using the ‘Measure’ function in FIJI. Puncta with mean intensity values under the identified threshold were excluded, and corresponding XY coordinates were exported for analysis. Resolution for each cell was quantified from the Full-Width Half Max (FWHM) across n=28-70 puncta from 2-3 2 µm x 2 µm regions in each image. Lineplots were used to measure the intensity profile across individual points, and gaussian curves were fit to these data [see github for Jupyter notebook] using the equation: 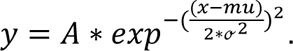

The FWHM was calculated from Gaussian fits to line intensity profiles of each punctum using the equation FWHM = ℴ*2.355 and averaged across the puncta for each image.

### Membrane Expression Quantification

Confocal images were also processed in FIIJI with a modified membrane quantification pipeline, as previously described^35^. In brief, a manually selected threshold for the GFP channel of each construct was used to segment individual cells. A 1.5 μm band was generated around each cell and the mean fluorescence intensity in this band for the 594 nm (Myc) channel was calculated using the ‘Measure’ function. Cells for which the segmentation pipeline failed to capture the membrane or segment individual cells were manually excluded. The mean fluorescence for each cell was normalized to the overall mean fluorescence for all day-matched wild-type Piezo1_Myc_ cells. In order to mimic conditions in which various Piezo constructs potentiate TREK1, and to control for varying levels of GFP expression in different vector backgrounds, we co-expressed each construct with TREK1. We found that the intensity of membrane fluorescence in the fly and mutant Myc constructs was equivalent to or slightly lower than wild-type mouse Piezo1_Myc_, indicating that increased expression of mutant constructs cannot explain their effects on TREK1 (**Figure S8D-E**). The image analysis pipeline for Figure S8 is available in the Grandl GitHub Repository: https://github.com/GrandlLab?tab=repositories.

### Spatial distribution analysis and modeling

Nearest Neighbor Distances (NNDs) between empirical TREK1 and Piezo1 coordinates were calculated using the KDtree function from the SciPy package in Python3^67^. To compare the empirical spatial relationship of TREK1 to Piezo1 puncta, we retained the empirical XY locations of Piezo1 puncta and simulated random populations of TREK1 puncta. Each simulation was performed 1,000 times per image. For random TREK1 distributions, for each image, TREK1 positions were simulated from a random distribution within the image ROI at a density equivalent to the empirical TREK1 density for that respective cell. Local Piezo1 densities were calculated for a 564 nm radius (=1 μm^2^ area) around each TREK1 punctum using the Ball Query function in SciPy.

Our estimate of 30 nm as the maximum cutoff for TREK1 and Piezo1 binding is based on the central location of the Piezo1 epitope, the predicted radius for Piezo1 in a biological membrane (14 nm^19,21,34^), the location of the TREK1 epitope and radius of the TREK1 channel (∼5 nm^68^), and some accounting for linkage error from our labeling strategy. Our estimate of 100 nm as the maximum cutoff for the Piezo1 footprint comes from estimates of the Piezo1 diameter (10-30 nm^20,23,25,26,34^ plus 5-fold the predicted decay length of the Piezo1 footprint in a biological membrane (14 nm)^24^.

Code and representative data for the images in **Figure 5** and **Figure 7** are available in the Grandl GitHub Server (2_Channel_Spatial_Analysis): https://github.com/GrandlLab?tab=repositories. Images, segmentation, and NND_T1-P1_ cumulative frequency distributions for all other cells are available on Dryad (https://datadryad.org/stash/share/8he-RNzsbsybFJmtKPP4puYIzl2j0wQI29iPdk_q19I).

