## Supplemental Figures for "Piezo1 ion channels are capable of conformational signaling"

Supplemental Figure 1

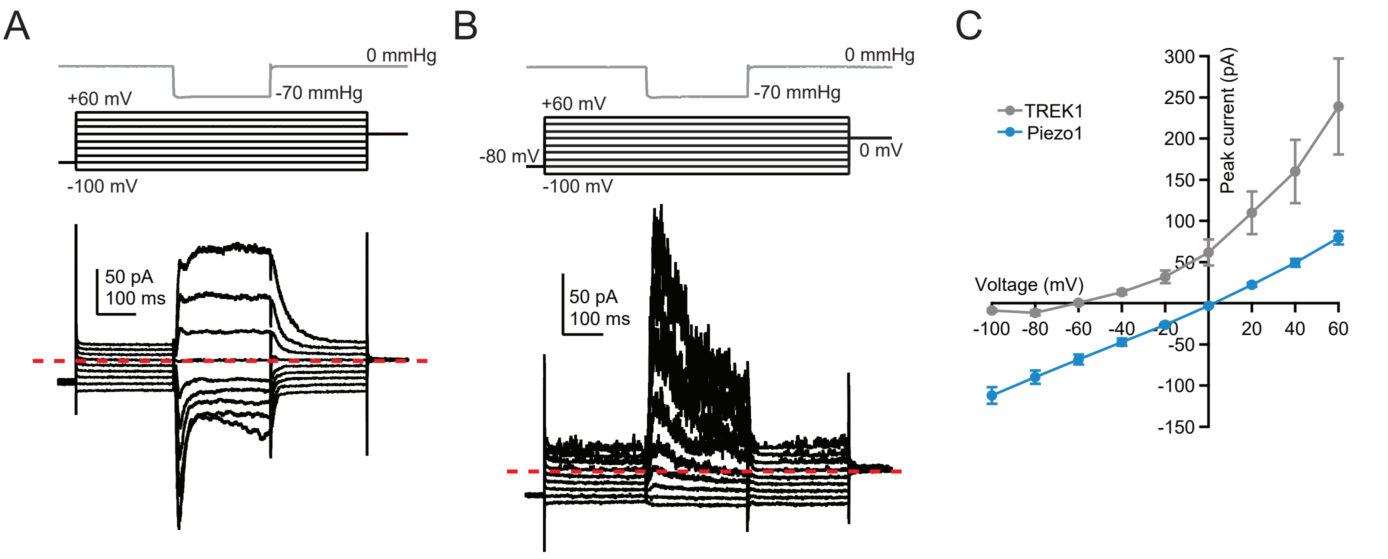

**Supplemental Figure 1. Reversal potentials for Piezo1- and TREK1-mediated currents.**

**A**, Pressure protocol (gray), voltage protocol (black), and representative current from a Neuro2A-Piezo1ko cell expressing mouse Piezo1. Pressure step was from 0 mmHg to -70 mmHg, voltages were from -100 mV to +60 mV in +20 mV increments. Red dashed line indicates zero current. **B**, as in A, for a cell expressing TREK1. **C**, Peak current-voltage relationships used to calculate reversal potential for cells expressing TREK1 (gray, 60.3±4.8 mV; n=8) or Piezo1 (1.4±0.8 mV, blue, n=17). Error bars are ±SEM.

Supplemental Figure 2

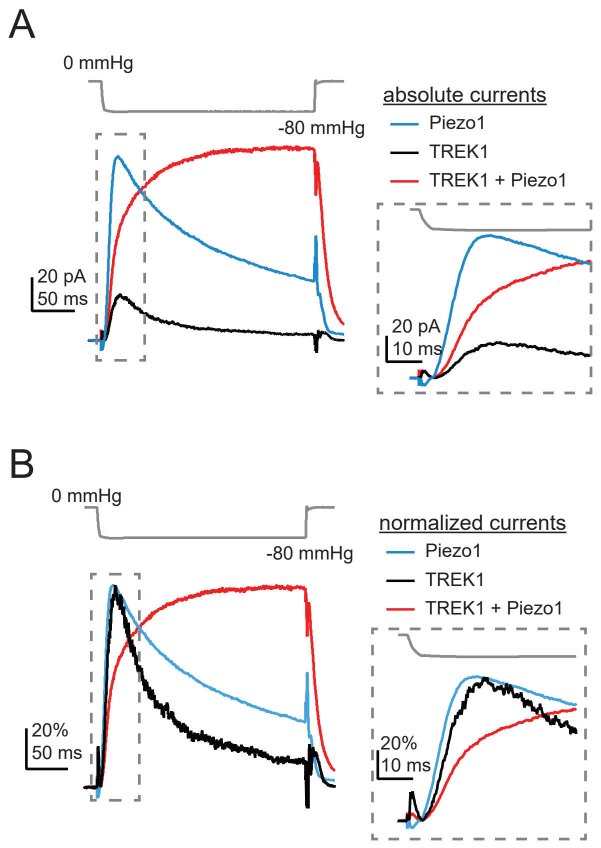

**Supplemental Figure 2. Apparent TREK1 activation kinetics are accelerated by inactivation in the absence of Piezo1.**

**A**, Pressure protocol (gray) and mean inward currents at -80 mV from N2A-Piezo1ko cells expressing Piezo1 (blue, n=25; current is inverted for display) and mean outward currents at 0 mV from cells expressing TREK1 (black, n=44), or Piezo1 and TREK1 (red, n=101). Pressure was -80 mmHg. Currents were baseline-subtracted, but not normalized prior to averaging. Inset, currents on a magnified time scale. Note that the inward Piezo1 current (blue) is fastest, followed by the outward currents through TREK1 in the presence of Piezo1 (red), and finally the outward currents through TREK1 in the absence of Piezo1 (black). **B**, Same currents as in A, normalized to their respective peaks. Note that the lack of inactivation of TREK1 currents in the presence of Piezo1, combined with the much larger peak amplitudes, leads to an apparent delayed rise time. Inset, currents on a magnified time scale.

### Supplemental Figure 3

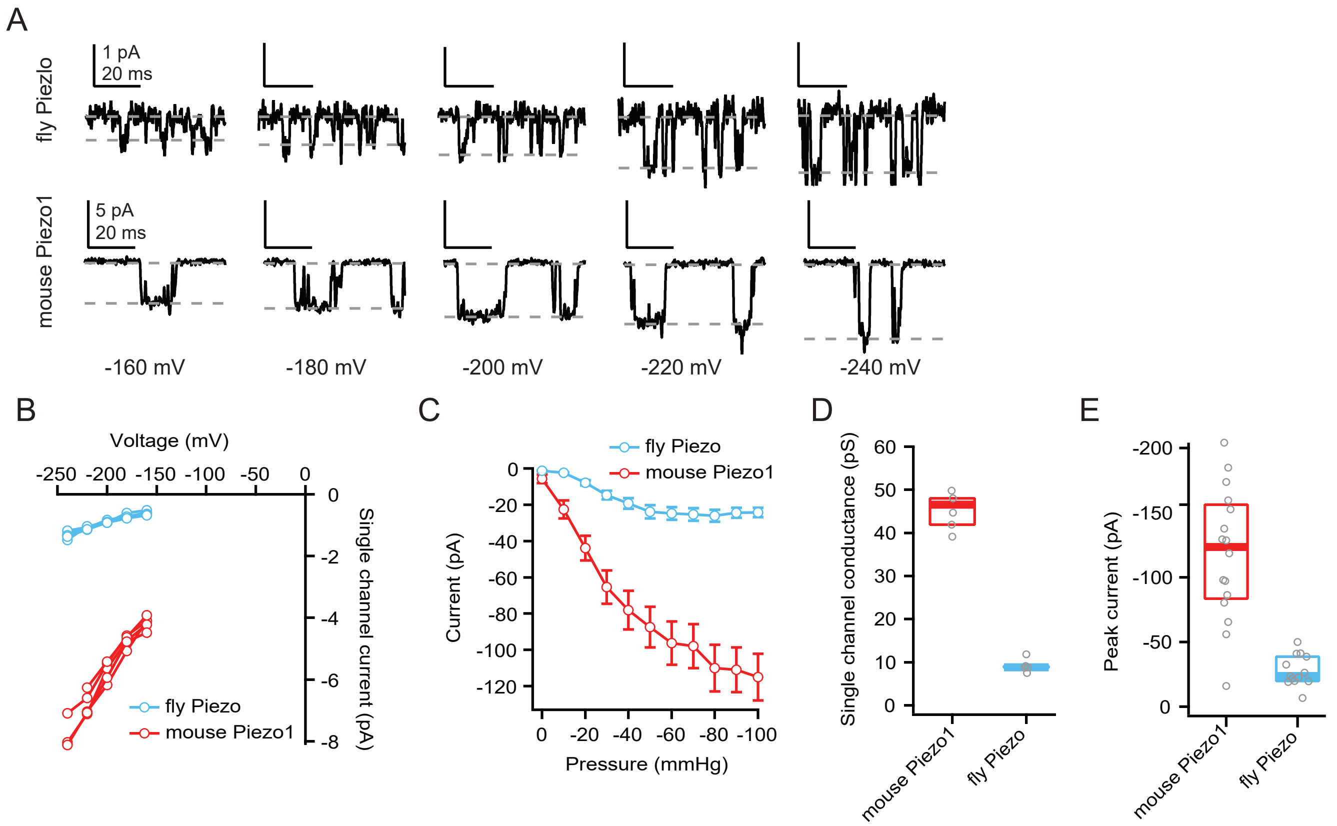

**Supplemental Figure 3. Slope conductance and macroscopic currents are ~5 fold smaller for fly Piezo compared to mouse Piezo1.**

**A**, Representative single-channel currents from Neuro2A-Piezo1ko cells expressing fly Piezo (top) or mouse Piezo1 (bottom) at -160 mV to -240 mV. Pressure was adjusted for each individual patch and voltage to elicit openings. **B**, Single-channel current as a function of voltage for each individual patch from cells expressing fly Piezo (n=5) and mouse Piezo1 (n=5). **C**, Peak current at -80 mV for patches expressing mouse Piezo1 (n=16) and fly Piezo (n=13). **D**, Slope conductance calculated from each individual patch in B. Median conductance was 44.7 pS for mouse and 8.8 pS for fly, yielding a ratio of 5.1. **E**, Box-and-whisker plots for peak inward current elicited by a pressure step to -80 mmHg at -80 mV. Median current was -124 pA for mouse and -24 pA for fly, yielding a ratio of 5.2.

Supplemental Figure 4

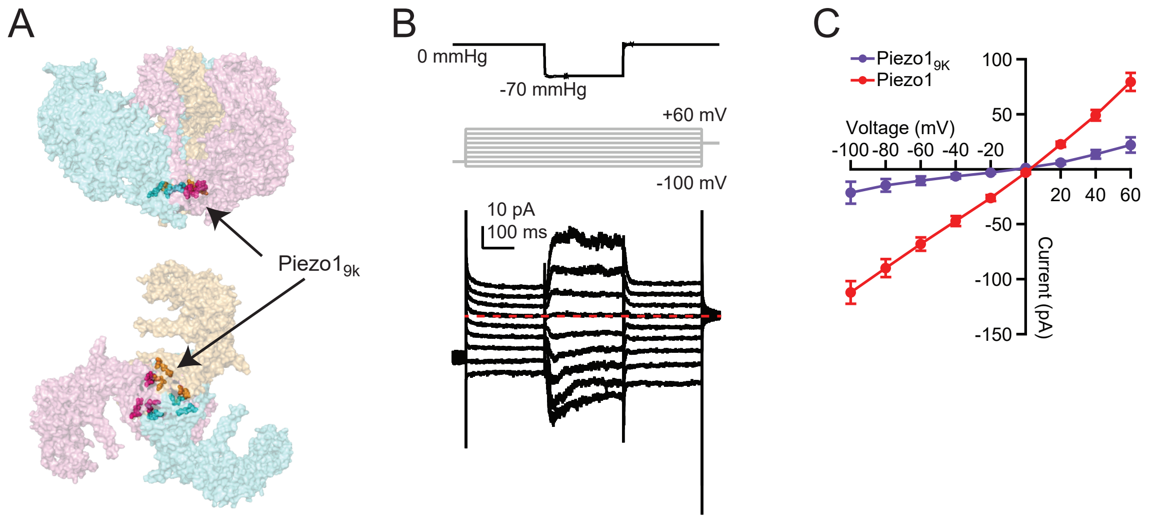

**Supplemental Figure 4. Characterization of Piezo1$_{9K}$ channel properties with pressure-clamp stimulation.**

**A**, Structural model of mouse Piezo1 (PDB: 6B3R[19]), highlighting residues mutated to lysine (Piezo1-SNCISESEE-9K[12]). Top, side-on view; bottom, bottom-up view. **B**, Pressure protocol (gray), voltage protocol (black), and representative currents (black) from a N2A-Piezo1ko cell expressing Piezo1$_{9K}$. Pressure steps were -70 mmHg, voltage steps were from -100 mV to +60 mV in +20 mV increments. Red dashed line indicates zero current. **C**, Mean peak current-voltage relationships for cells expressing wild-type Piezo1 (red, n=17) and Piezo1$_{9K}$ (purple, n=10).

Supplemental Figure 5

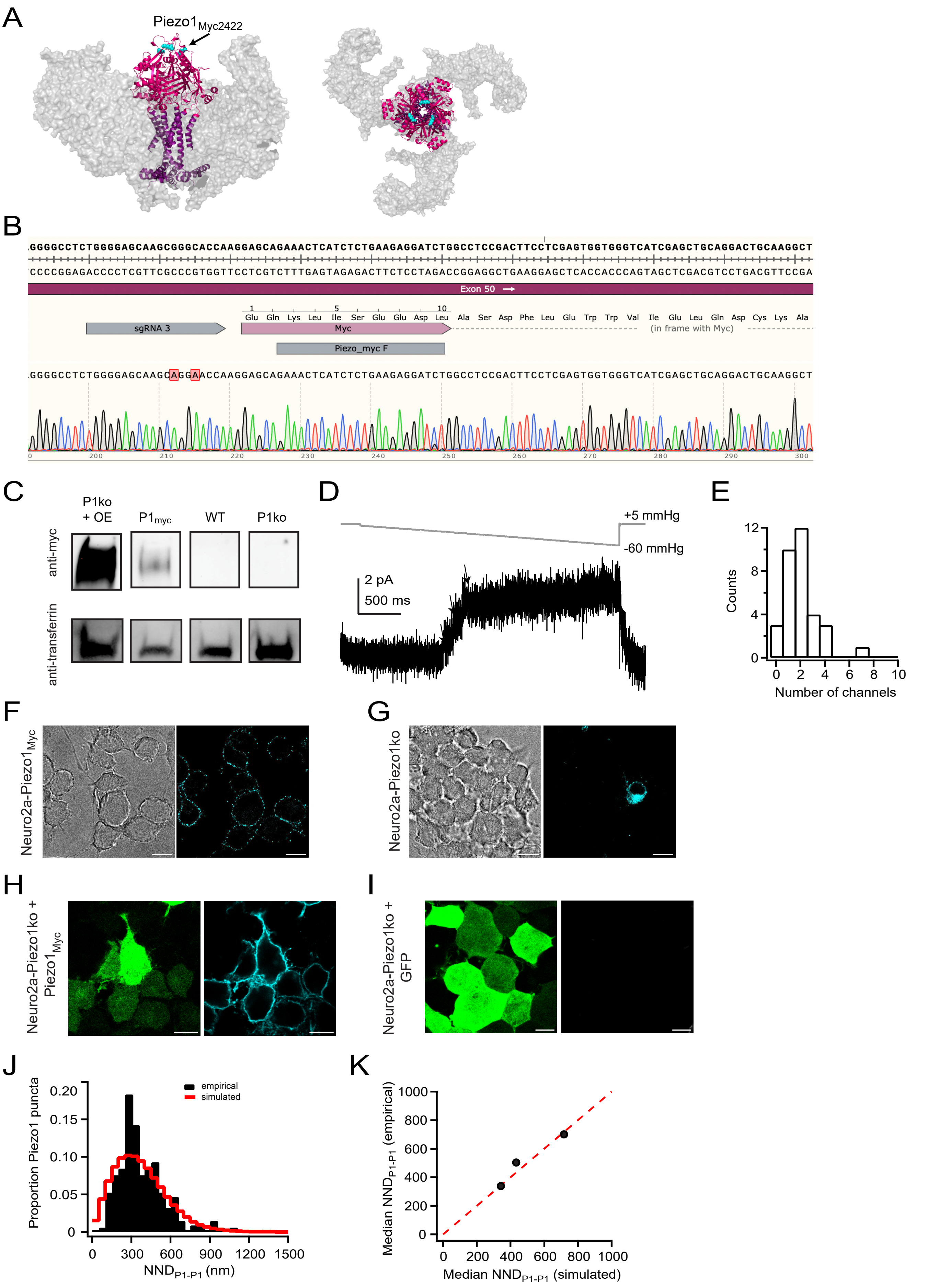

**Supplemental Figure 5. Validation and electrophysiological characterization of Neuro2a-Piezo1$_{MYC}$ cell line.**

**A**, Structural model of mouse Piezo1 residues 2192-2541 generated by AlphaFold3[69] (pore helices: purple, cap: magenta), superimposed on top of mouse Piezo1 blades (PDB: 6b3r[19]; gray), showing site of Myc tag insertion (cyan) following residue 2422 in an unstructured loop at the top of the cap domain. **B**, Strategy for insertion of a Myc tag immediately following residue 2422 in Piezo1 in Neuro2a cells. Two silent mutations (red) were introduced by the guide RNA to prevent re-cleavage by Cas9. Sequence chromatogram shows successful incorporation of the tag. **C**, Representative western blot showing membrane fraction isolated from Neuro2A-Piezo1ko cells overexpressing Piezo1$_{Myc2422}$ (P1ko + OE), Neuro2A-Piezo1$_{Myc}$ cells (P1$_{Myc}$), wild-type Neuro2A cells (WT), and Neuro2a-Piezo1ko cells (P1ko). Transferrin staining was used as a loading control. **D**, Pressure protocol (gray) and current (black) from a cell-attached patch with three channels from a Neuro2a-Piezo1$_{MYC}$ cell (arrows). Holding potential was +60 mV. **E**, Histogram of channel numbers in each patch. N=33 patches. **F**, Confocal images of Neuro2A-Piezo1$_{Myc}$ cells labelled with chicken anti-Myc antibody and Alexa Fluor 594 goat anti-chicken antibody. Brightfield (left) and 594 channel (cyan, right) are shown. Scale bar is 10 $\mu$m. **G**, As in F, for Neuro2A-Piezo1ko cells. 488 channel to visualize GFP (as a marker of transfection, green, left) and 594 channel (cyan, right) are shown. **H**, As in F, for Neuro2A-Piezo1ko cells overexpressing Piezo1$_{Myc}$. **I**, As in F, for Neuro2A-Piezo1ko cells overexpressing GFP. **I**, Black bars, normalized histogram of Piezo1-Piezo1 nearest neighbor distances (NND$_{P1-P1}$) for the cell shown in in **Figure 5A**. Red line, simulated NND$_{P1-P1}$ from a random distribution of equal density and area. **J**, Median NND$_{P1-P1}$ for empirical data as a function of NND$_{P1-P1}$ for simulated data. Dashed line represents unity. n=3 cells.

### Supplemental Figure 6

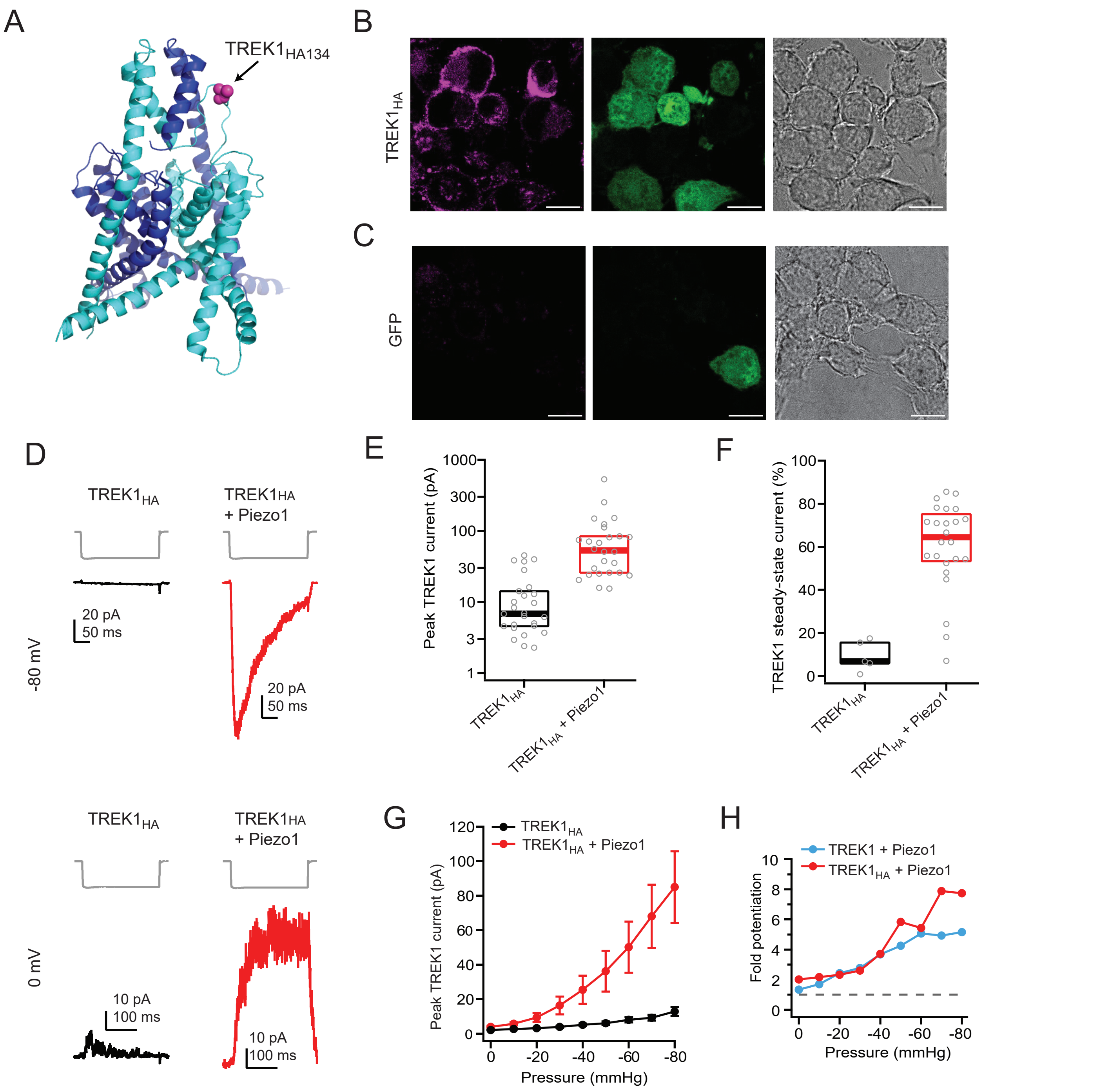

**Supplemental Figure 6. Validation and electrophysiological characterization of $TREK1_{HA}$ construct.**

**A**, Structural model of mouse TREK1 (PDB: 6CQ6[68]), highlighting the two monomers (light blue and dark blue) and the location of an incorporated HA tag at residue N134 (magenta) in the loop connecting the cap and the P1 pore helix in one monomer; this loop is not resolved in the other monomer. **B**, Confocal images of Neuro2A-$Piezo1_{Myc}$ cells expressing $TREK1_{HA}$ and labelled with rabbit anti-HA and ATTO 647N goat anti-rabbit. 647 channel (magenta, left), 488 channel to visualize GFP (as a marker of transfection, green, center) and brightfield (right) images are shown. Scale bar is 10 μm. **C**, as in B, for cells expressing GFP. **D**, Pressure protocol (gray) and currents from cell-attached patches from Neuro2A-Piezo1ko cells expressing $TREK1_{HA}$ (black), or $TREK1_{HA}$ + mouse Piezo1 (red), elicited by a 250 ms pressure step to -80 mmHg. Holding potential was either -80 mV (top) or 0 mV (bottom). **E**, Box-and-whisker plots for peak outward currents at 0 mV and -80 mmHg. n=26 ($TREK1_{HA}$), n=26 ($TREK1_{HA}$ + Piezo1). **F**, Box-and-whisker plots for TREK1 steady-state current at -80 mmHg. n=5 ($TREK1_{HA}$); n=24 ($TREK1_{HA}$ + Piezo1). **G**, Mean pressure-response curves at 0 mV, generated from full pressure protocols in A. **H**, Fold potentiation of TREK1 current by Piezo for wild-type TREK1 + Piezo1 (blue) and $TREK1_{HA}$ + Piezo1 (red), calculated as the ratio of median currents at each pressure ((TREK1 + Piezo1)/TREK1) and (($TREK1_{HA}$ + Piezo1)/$TREK1_{HA}$).

### Supplemental Figure 7

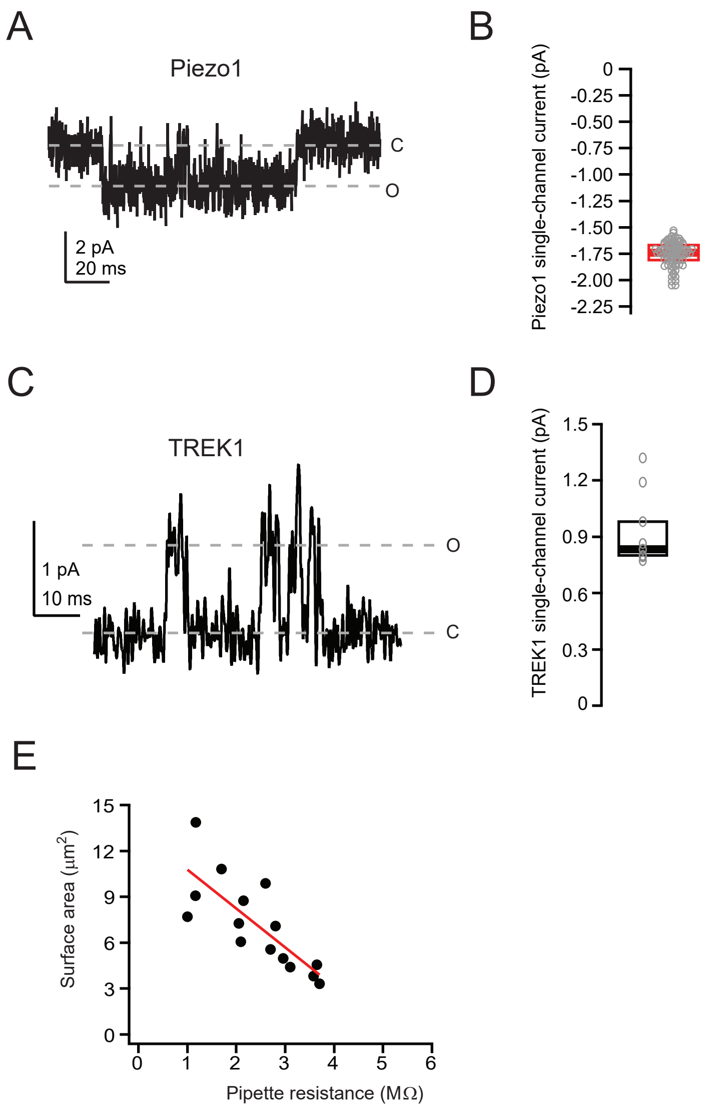

**Supplemental Figure 7. Measurement of TREK1 and Piezo1 single channel currents and calibration of pipette resistance and patch surface area.**

**A**, Representative single-channel opening from a Neuro2A-Piezo1ko cell expressing Piezo1. Voltage was -80 mV, pressure was 0 mmHg. **B**, Box-and-whisker plot for TREK1 single-channel current (n=87). **C**, Representative single-channel opening from a cell expressing TREK1. Voltage was 0 mV, pressure was -10 mmHg. Current was digitally filtered at 1 kHz for display. Dashed lines denote open (O) and closed (C) states. **D**, Box-and-whisker plot for TREK1 single-channel current (n=9). **E**, Surface area as a function of pipette resistance for cell-attached patches (data originally from [1]). Data were fit with a linear equation, $y = 13.33 – 2.55x$, which was then used to estimate patch dome surface area (y) for a given pipette resistance (x).

### Supplemental Figure 8

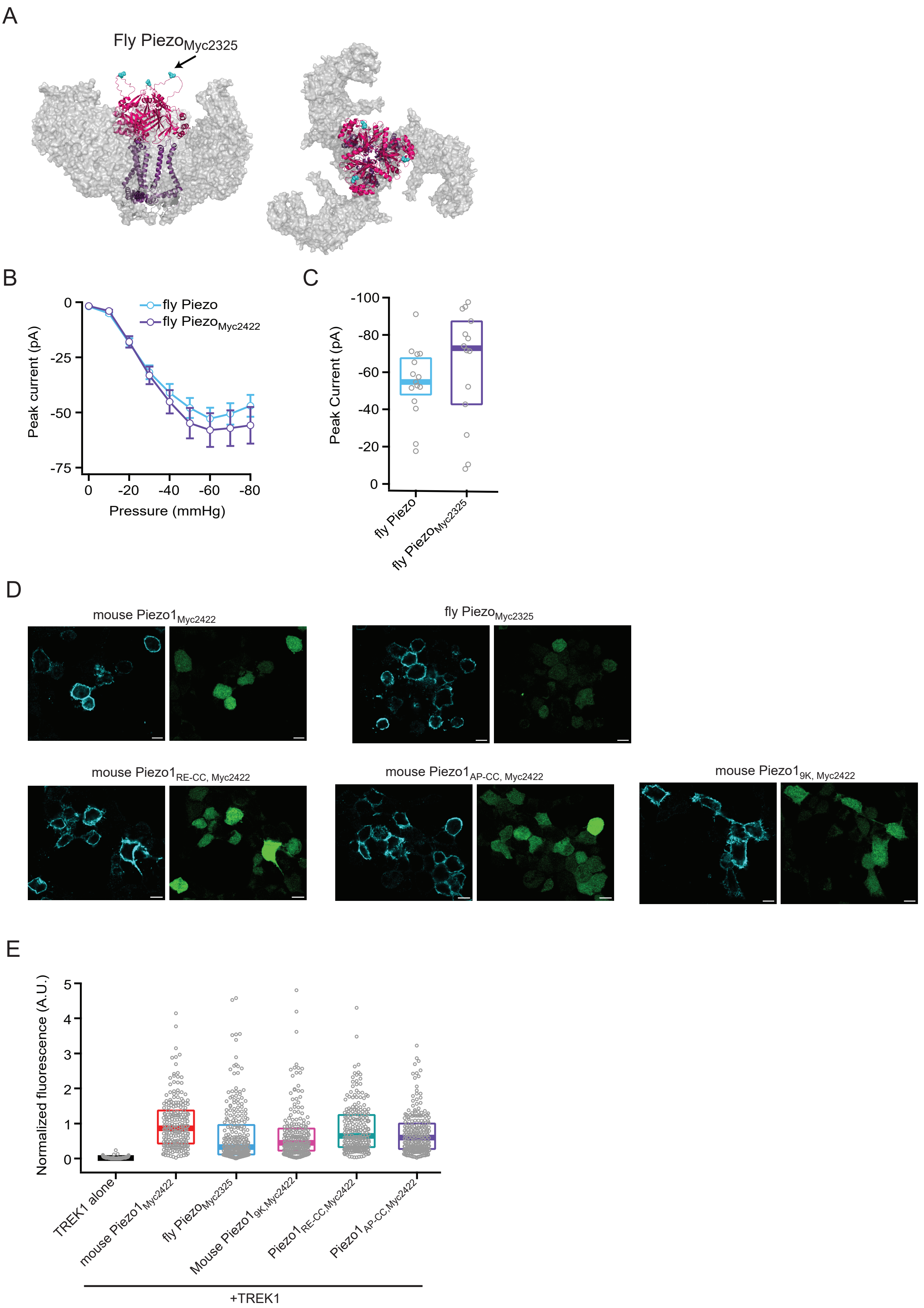

**Supplemental Figure 8. Membrane expression levels of Piezo constructs.**

**A**, Structural model of fly Piezo residues 2141-2515 generated by AlphaFold3[69] (pore helices: purple, cap: magenta), superimposed on top of mouse Piezo1 blades (PDB: 6b3r[19]; gray), showing site of Myc tag insertion (cyan) following residue 2325 in an unstructured loop at the top of the cap domain. **B**, Peak current at -80 mV as a function of pressure for patches expressing wild-type fly Piezo (n=15) and fly $\text{Piezo}_{\text{Myc2325}}$ (n=14). **C**, Box-and-whisker plots for peak inward current elicited by a pressure step to -80 mmHg at -80 mV for wild-type fly Piezo and fly $\text{Piezo}_{\text{Myc2325}}$. **D**, Confocal images of Neuro2A-Piezo1ko cells expressing TREK1 and the indicated $\text{Piezo}_{\text{Myc}}$ construct. Cells were labelled with chicken anti-Myc and AlexaFluor 594 plus goat anti-chicken. 594 channel (cyan, left) and 488 channel to visualize GFP (as a marker of transfection, green, center) are shown. Scale bar is 10 μm. **E**, Fluorescence intensity normalized to wild-type mouse $\text{Piezo}_{\text{Myc}}$ (A. U.) for cells transfected with TREK1 alone (median = 0.015, n=215 cells), TREK1 + mouse $\text{Piezo1}_{\text{Myc}}$ (median = 0.86, n=209 cells), TREK1 + fly $\text{Piezo1}_{\text{Myc}}$ (median = 0.33, n=244), TREK1 + mouse $\text{Piezo1}_{\text{AP-CC,Myc2422}}$ (median = 0.60, n = 216 cells), TREK1 + mouse $\text{Piezo1}_{\text{RE-CC,Myc2422}}$ (median = 0.64, n = 216 cells), and TREK1 + mouse $\text{Piezo1}_{\text{9K,Myc2422}}$ (median = 0.45, n = 209 cells).

**Supplemental Movie 1: Co-Labeled Piezo1$_{Myc}$ (Cyan) and TREK1$_{HA}$ (magenta).**

Movie shows the 100x, STED acquired z-stack of an entire Neuro2A-Piezo1ko cell from the bottom, (initial frames), to the cell surface (final frames).

| | | Channel Density (channels/μm²) | | Median Nearest-neighbor distance (nm) | | | |
|---|---|---|---|---|---|---|---|
| Cell ID | Image surface area (μm²) | TREK1 | Piezo1 | TREK1-Piezo1 | Piezo1-TREK1 | Piezo puncta with a TREK1 punctum within 30 nm (%) | Resolution (nm) |
| A | 178 | 1.9 | 1.9 | 290 | 295 | 1.1 | 88 |
| B | 61 | 2.3 | 1.3 | 346 | 239 | 2.6 | 96 |
| C | 354 | 1.3 | 0.4 | 657 | 352 | 0.6 | 80 |

**Supplementary Table 1. STED image properties for Neuro2A-Piezo1$_{Myc}$ cells transfected with TREK1$_{HA}$.**

Representative cell in Figure 5 is Cell “A”.

| | | Channel density (puncta/μm$^2$) | | Median nearest-neighbor distance (nm) | | |
|---|---|---|---|---|---|---|
| Cell ID | Image surface area (μm$^2$) | TREK1 | Piezo1 | TREK1-Piezo1 (empirical) | TREK1-Piezo1 (random model distributions) | Resolution (nm) |
| A | 183.9 | 4.9 | 4.6 | 169 | 212 | 91 |
| B | 28.4 | 1.9 | 7.7 | 132 | 156 | 66 |
| C | 138.3 | 2.9 | 4.3 | 190 | 218 | 66 |
| D | 112.2 | 4.0 | 2.2 | 300 | 348 | 96 |
| E | 67.0 | 2.8 | 3.3 | 189 | 258 | 76 |
| F | 118.5 | 2.5 | 7.1 | 99 | 165 | 75 |
| G | 179.3 | 4.1 | 5.5 | 153 | 198 | 77 |
| H | 138.3 | 1.9 | 3.8 | 174 | 225 | 79 |
| I | 146.3 | 4.4 | 2.9 | 231 | 275 | 98 |

**Supplementary Table 2. STED image properties for Neuro2A-Piezo1$_{ko}$ transfected with TREK1$_{HA}$ and Piezo1$_{Myc}$.**

Representative cell in Figure 7 is Cell “A”.
